# Inflammasome activation differences underpin different *Mycobacterium tuberculosis* infection outcomes

**DOI:** 10.1101/2025.06.06.658135

**Authors:** Ranjeet Kumar, Afsal Kolloli, Gunapati Bhargavi, Seema Husain, Saleena Ghanny, Patricia Soteropoulos, Selvakumar Subbian

**Affiliations:** The Public Health Research Institute at New Jersey Medical School, Rutgers University, Newark, NJ 07103, USA; Department of Microbiology, Biochemistry and Molecular Genetics, Rutgers, The State University of New Jersey, New Jersey Medical School, Newark, New Jersey, USA; Genomics Center, Rutgers, The State University of New Jersey, New Jersey Medical School, Newark, New Jersey, USA

## Abstract

The clinical outcome of *Mycobacterium tuberculosis* (Mtb) infection ranges from latent/non-progressive disease to active/progressive tuberculosis (TB), but the cellular events contributing to these variable outcomes remain unknown. Here, we report that progressive Mtb infection is associated with upregulation of guanylate-binding protein-1 (GBP1), hypoxia-inducible factor 1α (HIF-1α) and elevated NLR family pyrin domain-containing (NLRP3) inflammasome activation pathways. Using rabbit lungs and in primary rabbit and human macrophages as well as human THP-1 cell line-derived macrophages for infection with laboratory (H37Rv) or clinical Mtb strains (HN878 or CDC1551) that differ in virulence, we show that NLRP3 inflammasome activation by HIF-1α and GBP1 leads to elevated mitochondrial stress, apoptosis and necrosis during progressive infection by HN878. These biological functions and pathways are dampened in rabbit lungs, primary rabbit and human macrophages during non-progressive infection by CDC1551. These findings are consistent with and confirmed by Mtb infection studies of macrophages knocked-down for HIF-1α or GBP1 expression. Our study indicates that differences in HIF-1α- and GBP1-mediated NLRP3 inflammasome activation influence the outcome of Mtb infection to active or latent TB.

## 1. INTRODUCTION

Infection with *Mycobacterium tuberculosis* (Mtb) resulted in 10.8 million new cases of tuberculosis (TB) and 1.25 million deaths worldwide in 2023. Additionally, approximately a quarter of the global population is estimated to harbor asymptomatic latent Mtb infection (LTBI), which can reactivate to symptomatic TB upon immune-suppressing host conditions (WHO, 2024; Riccardi et al., 2020; Sotgiu et al., 2015). Diversity in pathogenic Mtb strains can differently modulate the host immune responses, leading to divergent outcomes of infection and TB transmission in humans (Chae and Shin, 2018; Comas and Gagneux, 2011; Kallenius et al., 2016; Smet et al., 2016; Stamm et al., 2015; Tientcheu et al., 2017). Similarly, the outcome of Mtb infection is impacted by the nature of infecting Mtb strains in the rabbit and non-human primate models of TB (Manabe et al., 2008; Peters et al., 2020; Zha et al., 2022). Unlike Mtb H37Rv, which is a standard laboratory strain, Mtb CDC1551 (CDC1551) and Mtb HN878 (HN878) are two well-characterized clinical Mtb strains, isolated from human TB outbreaks (Manabe et al., 2008; Manca et al., 2001; Tientcheu et al., 2017; Valway et al., 1998). Based on immune responses and disease pathologies elicited by these two strains in vitro and in animals, CDC1551 is deemed to be a “hyperimmunogenic” strain whereas HN878 is considered a “hypervirulent” strain (Merker et al., 2015; Hanekom et al., 2011; Koo et al., 2012; Manca et al., 1999; Reed et al., 2004; Subbian et al., 2011). Clinical and preclinical findings show that CDC1551 infection leads to a high rate of tuberculin skin test conversion in TB household contacts, elicits an early and effective pro-inflammatory cytokine response in monocytes, and induces a strong T cell activation in infected mouse lungs and established LTBI in rabbits (Manca et al., 1999; Manca et al., 2001; Subbian et al., 2012; Valway et al., 1998). In contrast, infection by HN878 is associated with a higher rate of active and drug-resistant TB cases and disease transmission in humans, does not mount host-protective innate and adaptive immune responses in mice, and causes the development of cavitary TB in the rabbit model (Merker et al., 2015; Hanekom et al., 2011; Barczak et al., 2005; de Jong et al., 2008; Manca et al., 2001; Subbian et al., 2011; Tsenova et al., 2005). In the rabbit model, we have shown that MtbHN878 infection leads to progressive disease in lungs, marked by necrotic and cavitary granulomas with high bacillary load (Subbian et al., 2011), whereas MtbCDC1551 infection results in a limited, non-progressive granulomatous response with protracted bacillary growth, which ultimately leads to the establishment of LTBI (Subbian et al., 2012). Moreover, in contrast to HN878-infected animals, the granulomas gradually resorbed and the bacterial load decreased in CDC1551-infected rabbits after 4 weeks of infection (Subbian et al., 2012; Subbian et al., 2011). Despite these observations, the difference in molecular and cellular pathways underpinning different host responses elicited by Mtb HN878 and CDC1551 are not fully understood.

Formation of granulomas, a highly organized cellular structure in the infected organs is a pathological hallmark of Mtb infection (Elkington et al., 2022; Kumar R and Subbian S. 2021). During progressive/active TB in humans, non-human primates, and rabbits, the elevated inflammatory response contributes to necrotic granulomas (NG) with hypoxic center and abundant bacterial presence (NG) (Kim et al., 2010; Manabe et al., 2008; Pena and Ho, 2015; Silva Miranda et al., 2012; Subbian et al., 2011). In contrast, non-progressive LTBI is marked by mostly fibrotic nodules (FN) with scanty bacterial presence and less inflammation (Kim et al., 2010; Manabe et al., 2008; Pena and Ho, 2015; Subbian et al., 2015). However, the cellular events underpinning NG and FN pathologies remains unknown.

Recent studies have highlighted a strong link between hypoxia and inflammation during Mtb infection in various host cells (Imtiyaz and Simon, 2010). The hypoxia-inducible factor-1 alpha (HIF-1α), a master regulator of immunologic and metabolic adaptations to hypoxia plays a key role in regulating inflammation, by promoting the expression of inflammatory molecules, including tumor necrosis factor α (TNFα), interleukin (IL) 6 and IL1β (Palazon et al., 2014). Importantly, expression of HIF-1α is upregulated in TB granulomas, and is associated with elevated proinflammatory responses (Denko, 2008; Li et al., 2021; Osada-Oka et al., 2019). One of the key pathways regulated by HIF-1α is activation of the inflammasome complex in myeloid cells and ablation of HIF-1α inhibits NLRP3 inflammasome activation (Huang et al., 2019).

Inflammasomes are intracellular sensory complexes activated in response to infection or cellular damage (Challagundla et al., 2022). These complexes comprise a sensor molecule, such as NLRP3 or absent in melanoma (AIM)2, the adaptor apoptosis-associated speck-like protein containing a CARD (ASC), and caspase (CASP)-1, and regulates cellular inflammatory responses (Nicholas et al., 2011). Activation of the inflammasome signaling pathway promotes elevated production of proinflammatory cytokines, such as IL1β, which contributes to host cell immunity during infection (Liu et al., 2016; Schroder and Tschopp, 2010; Zheng et al., 2020). Macrophages infected with Mtb activate NLRP3 and undergo death by multiple mechanisms, including necrosis (Escobar-Chavarria et al., 2023; Moreira et al., 2022; Ramon-Luing et al., 2022). Furthermore, NLRP3 inflammasome is activated by host-derived damage-associated molecular patterns (DAMPs), including mitochondrial reactive oxygen species (ROS) and DNA released during cellular stress (Bird, 2012; Shimada et al., 2012). Importantly, Mtb infection causes mitochondrial damage, alterations in mitochondrial mass, morphology, bioenergetic metabolism, and elevated cellular ROS production in macrophages, which subsequently results in inflammasome activation (Cumming et al., 2018; Patrick and Watson, 2021). Virulent Mtb has been reported to avert inflammasome activation to survive intracellularly (Rastogi and Briken, 2022; Van Hauwermeiren and Lamkanfi, 2023) However, regulation of inflammasome activation by divergent Mtb strains in macrophages, and the link between inflammasome activation and progressive versus non-progressive Mtb infection in the lungs are not fully understood.

Guanylate-binding proteins (GBPs) constitute a family of host proteins induced by both type I and type II interferons (IFNs) in cell culture and in vivo models (Honkala et al., 2019; Kim et al., 2011; Man et al., 2017; Pilla-Moffett et al., 2016; Shenoy et al., 2012). GBPs facilitate the destruction of microbial pathogens and liberating antigens that trigger innate immunity and activate inflammasomes in macrophages and monocytes (Man et al., 2017; Ngo and Man, 2017). Although prior studies suggest a causal link between inflammasome activation and the expression of HIF-1α and GBPs in regulating cellular immune response during TB, the differential regulation of inflammasome activation pathways in macrophages upon infection with various clinical Mtb strains has not been explored.

In the present study, we report different regulations of the inflammasome activation in the context of GBP1 and HIF-1α signaling pathways in the lungs and bone marrow-derived macrophages (rBMDM) of rabbits and in human macrophages infected with CDC1551 or HN878. We further elaborated our study to identify host effector immune mechanisms particularly, cell death pathways modulated by GBP1 and Hif-1α, in macrophages infected with different strains of Mtb. Our observations suggest that Mtb strain-dependent differences in regulating the inflammasome activation by GBPs and HIF-1α contributes to the difference in the pulmonary disease pathology between HN878 and CDC1551 infections in vivo and in vitro. These findings will aid in a better understanding of TB pathogenesis and guide the development of novel host-directed therapeutics for effective TB control.

## 2. RESULTS

### 2.1. Differential lung disease pathology in progressive and non-progressive Mtb infection

We first compared the gross pathology, histology and genome-wide transcriptome of rabbit lungs infected with HN878 or CDC1551 at the peak bacterial burden stage (i.e, 4 weeks post-infection). Several clearly visible subpleural granulomas were noted in the lungs of HN878-infected rabbits (Figure 1A) and the histological analysis showed large, highly cellular granulomas with inflammation, central necrosis (Figure 1B) and the presence of abundant neutrophils (Figure 1C). In contrast, no gross subpleural granulomas were seen in CDC1551-infected rabbit lungs (Figure 1D) and the histologic examination revealed few, small granulomas without remarkable necrosis or abundant presence of neutrophils in these lungs (Figure 1E and F). These findings are consistent with our previous study (Subbian et al., 2012). The elevated disease pathology in HN878-infected rabbit lungs compared to CDC1551-infected rabbit lungs correlated with a significantly higher bacillary load (Figure 1G). To determine whether these two Mtb strains that produce progressive versus non-progressive lung diseases differ in their intracellular growth in macrophages, we infected rabbit bone marrow derived macrophages (rBMDMs), human peripheral blood monocyte-derived macrophages (hu-MΦ) and human monocytic THP-1-derived macrophages (THP-1) with HN878 or CDC1551 or H37Rv (a standard laboratory strain) and determined the bacterial load up to 96 hours post-infection (hpi). We observed a significantly higher bacterial load at 96 hpi in rBMDMs, hu-MΦ and THP-1 infected with HN878, compared to CDC1551, and H37Rv at 96 hpi (Supplementary Figures 1A-C). Although H37Rv grew slightly better than CDC1551 in primary macrophages, the difference was not statistically significant. This observation indicates Mtb strain-specific growth in macrophages and highlights the similarity between rBMDMs and hu-MΦ in their capacity to control Mtb infection.

**Figure 1.**
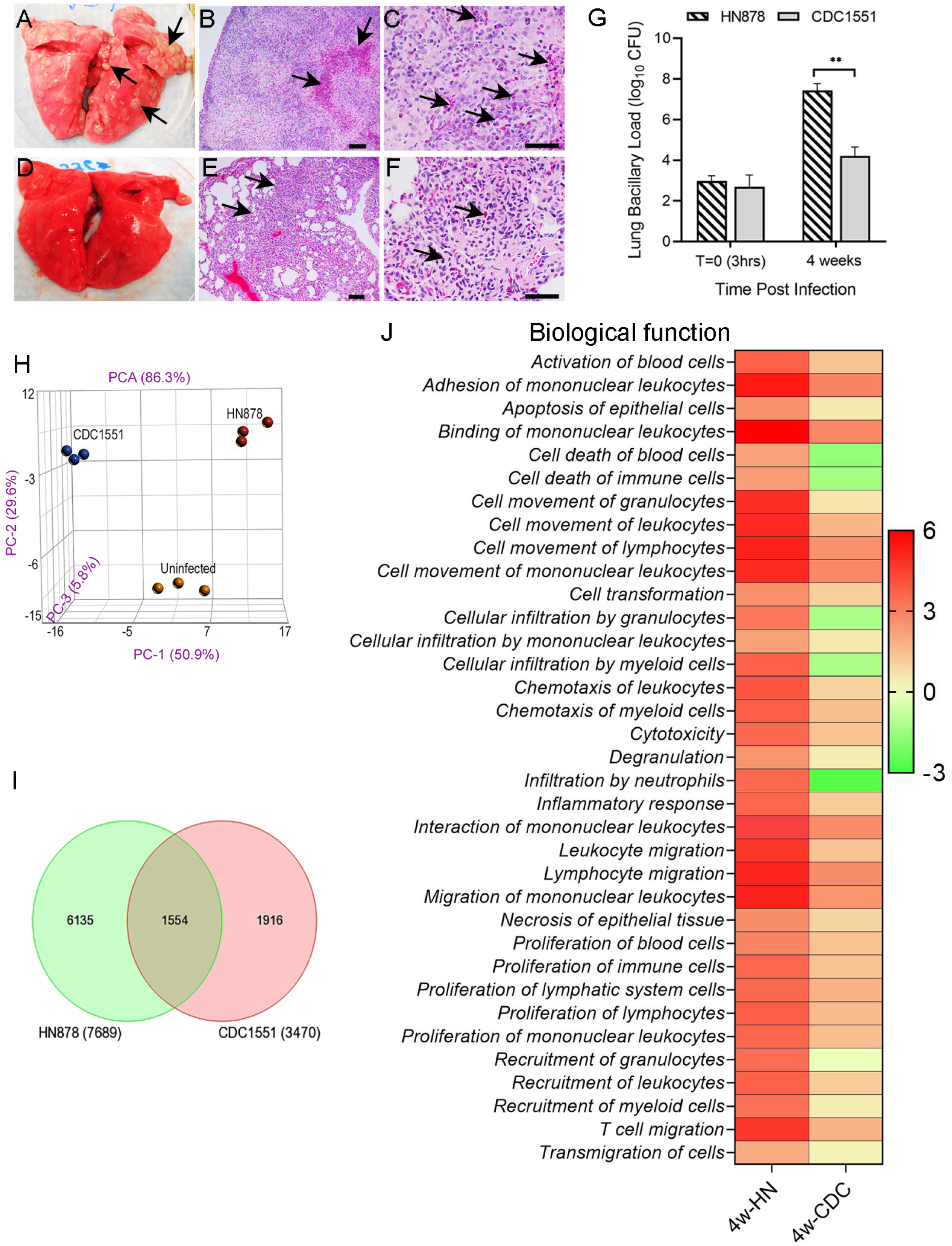
Different pathologies, bacterial loads and genome-wide transcriptomes of rabbit lungs infected with Mtb HN878 and CDC1551. (**A**) Gross pathology of rabbit lungs 4 weeks after infection with HN878 with clearly visible subpleural granulomas (arrows in A). (**B**) A representative low-magnification histologic image showing a big and highly cellular granuloma with prominent central necrosis (arrows). (**C**) A high-magnification histologic image showing a granuloma with extensive neutrophil infiltration near the necrotic region (arrows). (**D**) Gross pathology of rabbit lungs 4 weeks after infection with CDC1551 without any visible subpleural granuloma. (**E**) A representative low-magnification histologic image of rabbit lungs infected with CDC1551 for 4 weeks showing small and highly cellular granuloma (arrows). (**F**) A representative high-magnification histologic image of rabbit lungs infected with CDC1551 for 4 weeks showing granuloma with mild neutrophil infiltration (arrows). Scale bars in **B** and **E** represent 100µm and those in **C** and **F** represent 50 µm. (**G**) Lung bacillary load immediately after infection (T=0; 3hrs) and after 4 weeks of infection with Mtb HN878 or CDC1551. Values plotted are mean +/-SD of the number of bacterial CFU (n=5 per group per time point). **P<0.01 by Student t-test. (**H**) Principal component analysis (PCA) plot of the genome-wide transcriptomes of uninfected, HN878- or CDC1551-infected rabbit lungs at 4 weeks post-infection (n=3 per group). (**I**) Venn diagram of SDEG in rabbit lungs 4 weeks after infection with HN878 (green) or CDC1551 (red), compared to the uninfected controls (n=3 per group). (**J**) Heat map of top biological functions enriched in HN878 (HN)- or CDC1551 (CDC)-infected rabbit lungs at 4 weeks post-infection, compared to uninfected controls. Values plotted and shown in the scale bar are z-scores. Red color indicates upregulation, and green color indicates downregulation of specific pathways, compared to the uninfected controls (n=3 per group).

### 2.2. Progressive Mtb infection activates a robust inflammatory response network in the lung

To determine the immunological networks of progressive versus non-progressive Mtb infection in the lungs, we performed a comparative analysis of the microarray-based genomewide transcriptome profile of rabbit lungs infected with Mtb HN878 or CDC1551 for four weeks (GEO-GSE33094; GSE39219). Although these datasets were published previously, a direct comparison analysis between the datasets was not reported earlier (Subbian et al., 2012; Subbian et al., 2011). The principal component analysis (PCA) of CDC1551- or HN878-infected rabbit lung transcriptomes showed clear segregation to each other and to the uninfected group (Figure 1H). The number of significantly differentially expressed genes (SDEGs), determined by a cut-off p-value of <0.05 between Mtb-infected and -uninfected groups, showed 2.2-fold more SDEGs in the HN878-infected lungs than in the CDC1551-infected lungs (7,689 versus 3,470 genes) (Figure 1I). Further, the number of unique SDEGs in the HN878-infected group was 3.2 folds more than that of the CDC1551-infected group (6,135 genes versus 1,916 genes). Among all SDEGs, 1,554 were shared between HN878- and CDC1551-infected lungs.

The gene ontology (GO) analysis revealed that activation, migration, stimulation, proliferation, and necrosis of immune cells as well as the inflammatory response were highly upregulated in HN878-infected lungs, as shown by the z-score of significance (Figure 1J). In the CDC1551-infected lungs, cellular development, movement, growth, proliferation, maintenance, and homeostasis were the top-upregulated GO functions, whereas functions associated with cell death and neutrophil recruitment were significantly downregulated in this group (Figure 1J). Consistent with the GO analysis, distinct canonical immune pathways were perturbed between HN878- and CDC1551-infected rabbit lungs (Supplementary Figure 1D). Several inflammatory response pathways, including cytokine storm signaling, acute phase response, cell death signaling, and host cell destruction, were upregulated in the HN878-infected rabbit lungs (Supplementary Figure 1D). Neutrophil degranulation and neutrophil extracellular trap signaling pathways were also upregulated, consistent with the elevated neutrophil presence, in the HN878-infected lungs (Figure 1C and 1J). These inflammatory signaling pathways were downregulated, while eNOS and cAMP-mediated signaling were upregulated in the lungs of CDC1551-infected animals (Figure 1J and Supplementary Figure 1D).

We have previously reported that necrotic granulomas (NG) in human TB lungs were highly inflammatory and enriched with acid-fast bacilli (AFB+), compared to the fibrotic nodules (FN) that had dampened inflammation and mostly devoid of Mtb (GEO-GSE20050; Subbian et al., 2015). Consistent with these differences in disease presentation, we observed distinct immune network gene expression profile between NG and FN (Supplementary Figure 2A). These observations in human lung TB lesions further validate our findings in the rabbit models of TB. Particularly, expression of inflammatory genes such as *IL6*, *TNFA*, *IL21*, *CXCL9*, *CXCL10*, *CCL2*, *CCL5*, *NLRP3*, *STAT1*, *TBX21*, *CASP1* and *CASP3* were significantly upregulated in human NG and HN878-infected rabbit lungs, compared to the human FN and CDC1551-infected rabbit lungs, respectively (Supplementary Figures 2A-C and 3).

### 2.3. NLRP3 Inflammasome activation in progressive versus non-progressive Mtb lungs infections

To determine whether progressive and non-progressive Mtb infections differently regulate the NLRP3 inflammasome activation, we interrogated corresponding gene expression profile of human lung NG and FN, and rabbit lungs infected with HN878 or CDC1551. We observed significantly upregulated expression of *NLRP2*, *NLRP3*, *NLRP4*, *ASC*, *IL1B*, *CASP1*, *CASP8*, *TLR4*, *MYD88*, *CTSB* and *NEK7* in human lung NG and HN878-infected rabbit lungs. Of these genes, expression of *NLRP2*, *NLRP3*, and *ASC* were also upregulated in human lung FN and CDC1551-infected rabbit lungs, although to a lesser extent, compared to human lung NG and HN878-infected rabbit lungs, respectively (Figure 2A, B and Supplementary Figure 2B). Subtle differences between the non-progressive human FN and CDC1551-infected rabbit lungs were also noticed. For example, expressions of *NLRP4* and *TLR4* were downregulated in the human lung FN, while these genes were upregulated in CDC1551-infected rabbit lungs. Similarly, *CASP1, CASP8* and *MYD88* were not significantly expressed in the CDC1551-infected rabbit lungs, while these genes were mildly upregulated in human-lung FN, compared to the uninfected controls (Figure 2B and Supplementary Figure 2B). These differences could be inherent to the samples used, human lung FN versus rabbit CDC1551-infected diffused/healing type granulomas in lungs, although both were representative of non-progressive Mtb infection. Together, these results indicate that progressive Mtb infection is associated with a stronger NLRP3 inflammasome activation.

**Figure 2.**
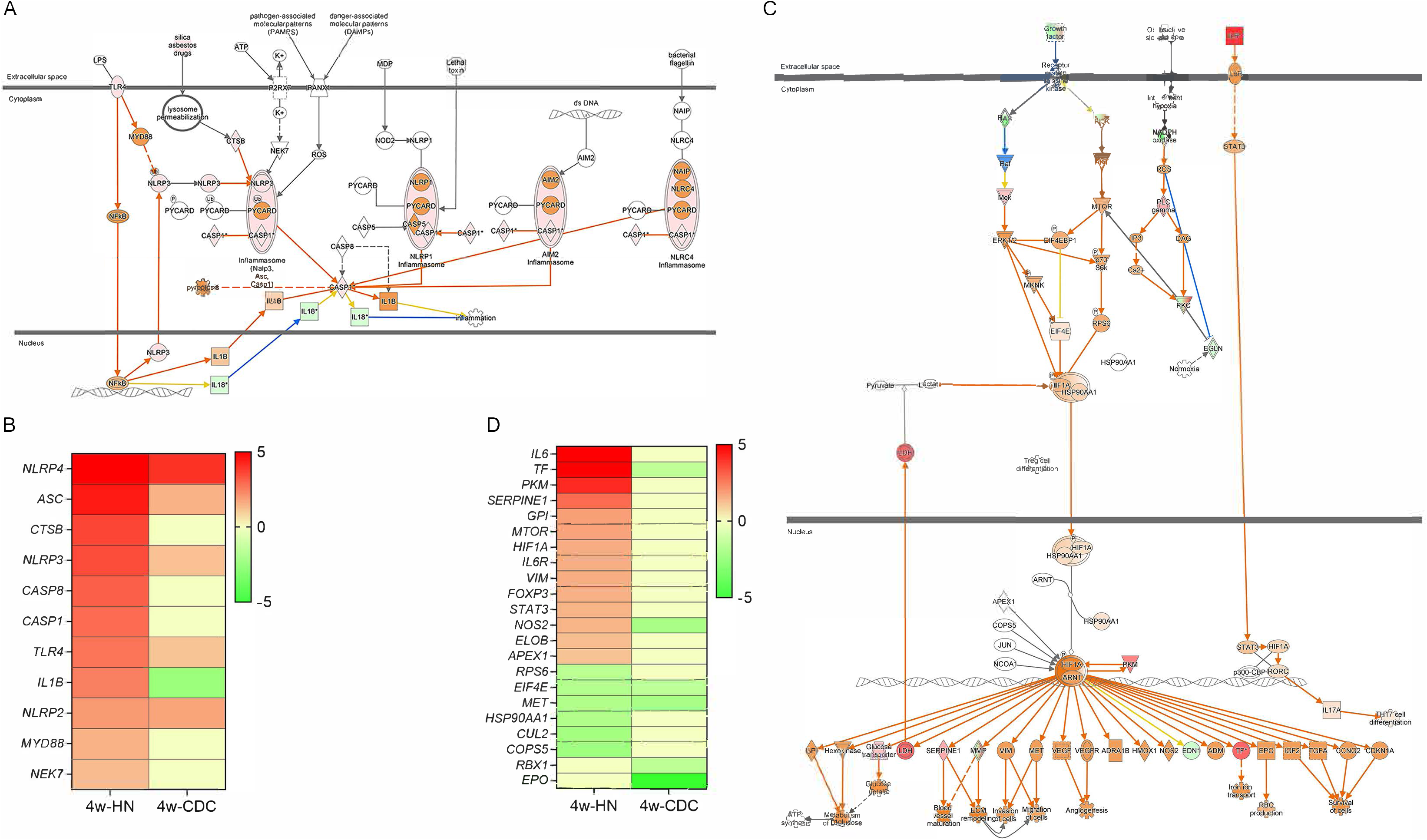
Differential expression of inflammasome activation and HIF-1α signaling pathways in rabbit lungs infected with Mtb HN878 or CDC1551. (**A**) Inflammasome activation pathway map showing interactions among member genes. (**B**) Heat map showing the expression of the inflammasome activation pathway member genes in HN878 (HN)- or CDC1551 (CDC)-infected rabbit lungs. (**C**) HIF-1α signaling pathway map showing interactions among member genes. (**D**) Heat map showing the expression of the HIF-1α signaling pathway member genes in HN- or CDC-infected rabbit lungs. In **A** and **C**, the expression level in rabbit lungs infected with HN878 at 4 weeks, relative to uninfected controls, was used on the maps. The red color indicates upregulation, and the green color indicates downregulation. The orange line indicates positive regulation, and the blue line indicates negative regulation among member genes. In **B** and **D**, the red color indicates upregulation, the green color indicates downregulation, and the yellow color indicates no significant change.

### 2.4. Expression of HIF-1α and NLRP3 are upregulated during progressive Mtb infection in the lungs

Since HIF1α has been shown to be associated with inflammatory response and NLRP3 inflammasome activation networks, which were differently regulated between progressive and non-progressive Mtb infection, we tested the expression pattern of HIF-1α signaling network genes in these two conditions in human and rabbit lungs. Expressions of several HIF-1α signaling network genes, including *HIF1A*, *IL6*, *TF*, *PKM*, *SERPINE1*, *MTOR*, *GPI*, *STAT3, APEX1, VIM* and *NOS2* were significantly upregulated in human NG and HN878-infected rabbit lungs, whereas they were downregulated in human FN and CDC1551-infected rabbit lungs (Figure 2C and D and Supplementary Figure 2C). Consistent with this transcriptome profiles, spatial expression levels of HIF-1α and NLRP3 were significantly upregulated in HN878-infected rabbit lungs, compared to CDC1551-infected and uninfected rabbit lungs (Figure 3A-D). Interestingly, spatial expression of NLRP3 was also significantly upregulated in CDC1551-infected rabbit lungs, compared to the uninfected counterpart, although the levels were about three times lesser than that observed in HN878-infected lungs (Figure 3D).

**Figure 3.**
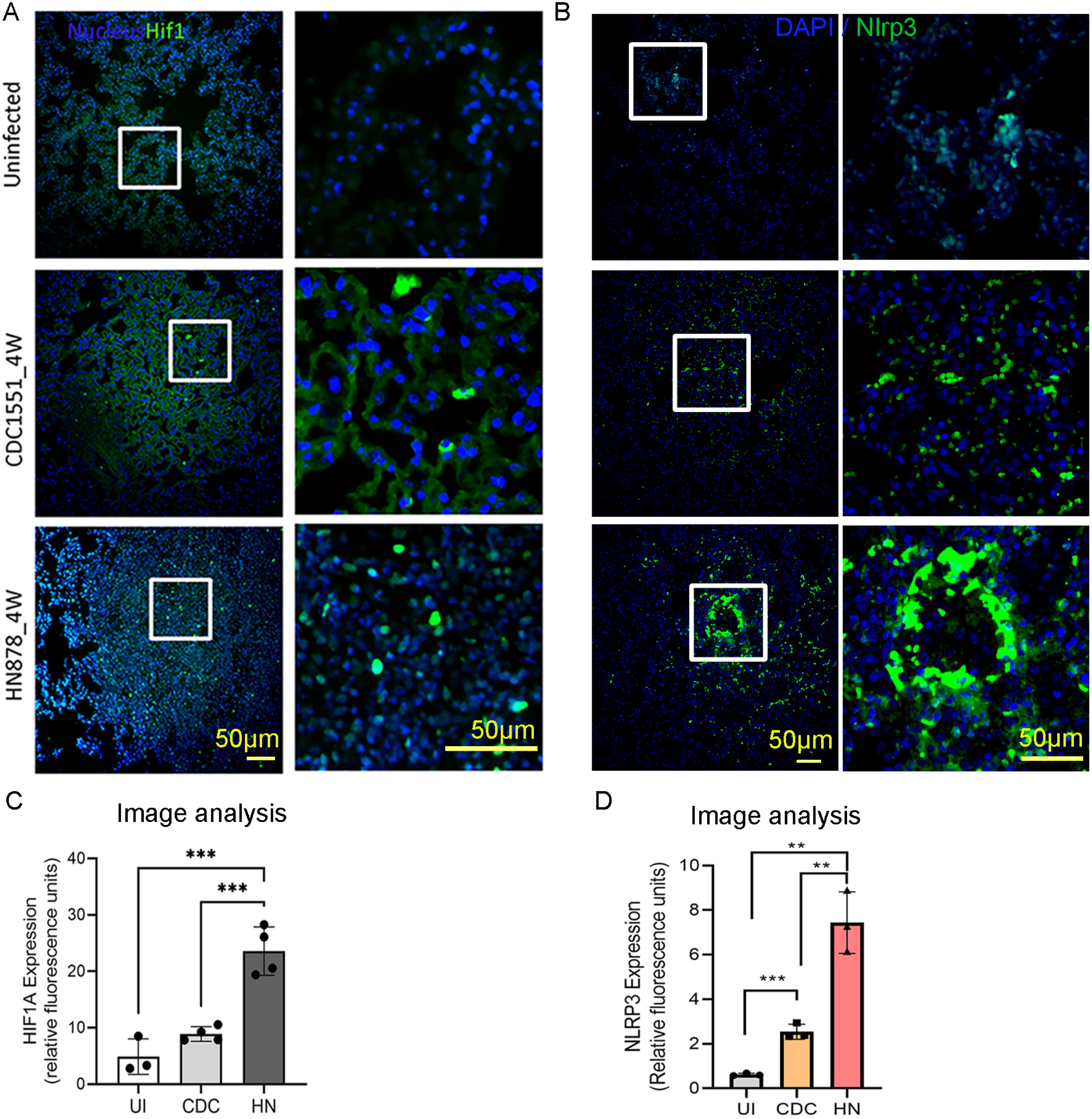
Spatial expression of HIF-1α and NLRP3 in Mtb HN878- or CDC1551-infected rabbit lungs. Representative images (**A**) and quantification (**C**) of HIF-1α in rabbit lung section with or without CDC1551 or HN878 infection at 4 weeks post-infection probed with a fluorescently labeled HIF-1α antibody. **(B and D)** Representative images (**B**) and quantification (**D**) of NLRP3 in rabbit lung section with or without CDC1551 or HN878 infection at 4 weeks post-infection probed with a NLRP3 antibody, as mentioned in the method section. Right images in **A** and **B** are magnified boxed areas in corresponding left images to show the labeling of cells (green signals). Experiments are representative with n=3-4 rabbits in each group, and statistical analyses were performed using one-way ANOVA. *P<0.05; ***P<0.005; ****P<0.001.

### 2.5. Differential activation of inflammasome and HIF-1α signaling pathways in macrophages infected with different Mtb strains

Previous studies have shown that HIF-1α can activate the NLRP3 inflammasome in human macrophages (Nicholas et al., 2011). Because we observed significant activation of both HIF1α and inflammasome activation pathways in rabbit lungs infected with HN878, compared to CDC1551 (Figures 2 and 3), we investigated whether macrophages infected with HN878 and CDC1551 would regulate the expression of these pathway genes differently.

Using real-time qPCR, we analyzed the expression profiles of *HIF1A*, together with selected NLRP3-inflammasome complex genes (*NLRP3* and *ASC*) and inflammatory effector molecules (*IL1B*, *IL6* and *TNFA*) in rabbit lungs, rBMDMs, THP-1 and hu-MΦ infected with HN878 or CDC 1551 or the laboratory Mtb strain H37Rv. We found significantly increased expression of *HIF1A*, *NLRP3*, *ASC, IL1B*, *IL6* and *TNFA* in HN878-infected rabbit lungs, rBMDM, THP-1 and hu-MΦ, compared to those infected with CDC1551 (Figure 4A-D). These observations suggest significant activation of HIF1A, components of NLRP3 inflammasome and its effector molecules during active/progressive Mtb infection and highlight the similarities in the host responses among rabbit lungs, rBMDM, and hu-MΦ following HN878 or CDC 1551 infection. The expression profile of many of the tested genes in H37Rv infected macrophages were like that of CDC551-infected human cells, while ASC and IL6 were not significantly expressed in rBMDMs infected with H27Rv (Supplementary Figure 4).

**Figure 4.**
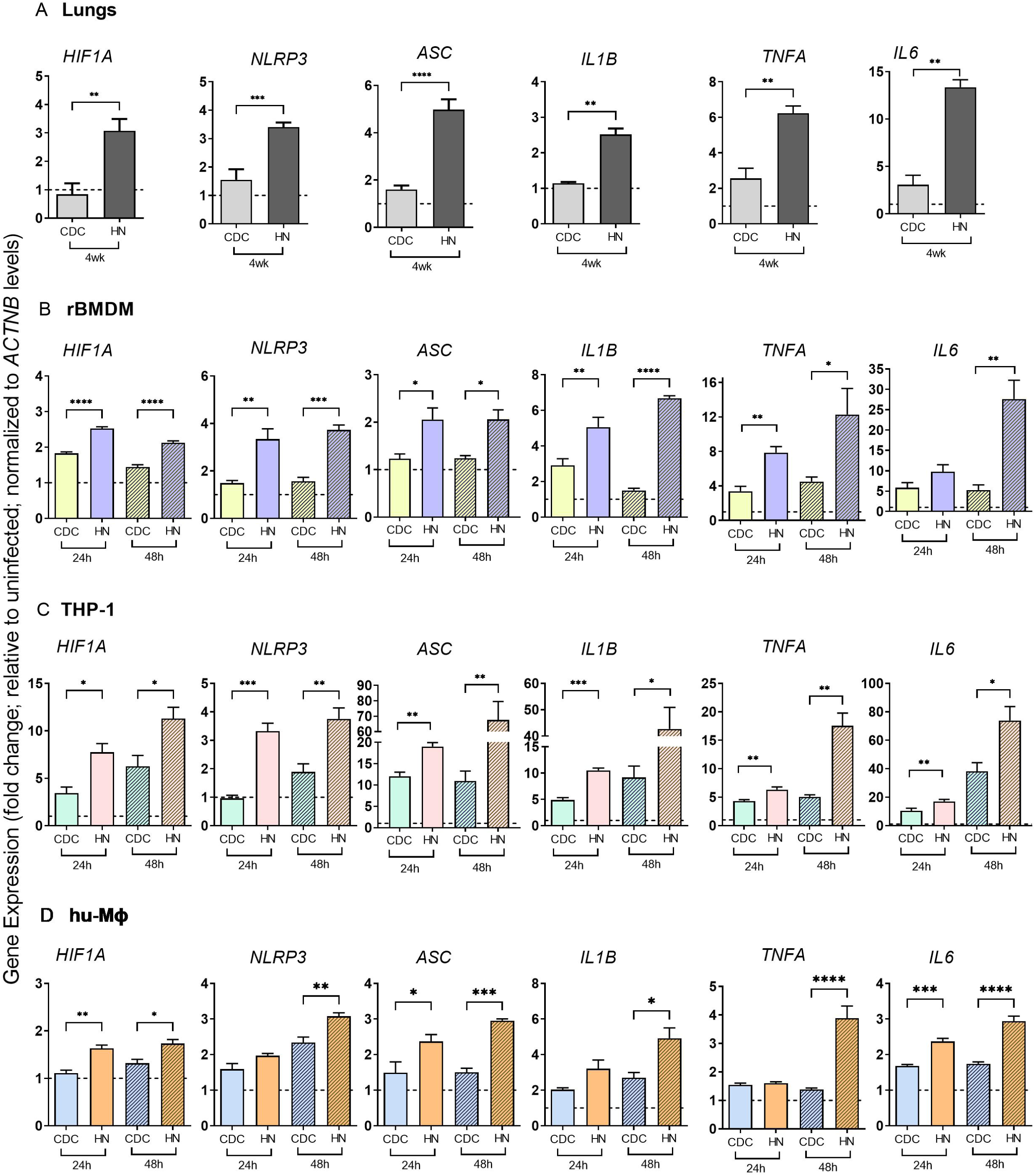
Expression profile of genes in the NLRP3 inflammasome activation pathway during Mtb HN878 or CDC1551 infection. qPCR was used to measure the transcript levels of *HIF1A*, *NLRP3*, *ASC*, *IL1B*, *TNFA* and *IL6* in CDC1551 (CDC)- or HN878 (HN)-infected rabbit lungs 4 weeks post-infection (**A**) or in rBMDMs (**B**) THP-1 (**C**) or human PBMC-derived macrophages (hu-MΦ) infected with CDC or HN for 24 or 48 hours. Fold changes in expression levels of indicated genes in Mtb-infected samples relative to those in uninfected samples (dotted lines) are calculated. The level of *ACTNB* expression is used to normalize the expression level of test genes. (Data are representative of three independent experiments performed with n=3-5 samples per group. Statistical analyses were performed using Student’s *t* -test. *P<0.05; ** P < 0.01; ***P<0.005.

To further confirm and validate differences in the NLRP3 inflammasome activation pathway in macrophages at the protein level, we performed a Western blot analysis of THP-1 macrophages infected with HN878 or CDC 1551 (Figure 5A). Consistent with the mRNA analysis, the Western blot analysis showed elevated levels of NLRP3, ASC, IL1β (precursor and mature forms), and Caspase-1 (precursor and cleaved forms) in Mtb-infected hu-MΦ as early as 4 hpi, compared to the uninfected controls. Moreover, compared to CDC1551-infected macrophages, the levels of NLRP3, ASC, IL1β and Caspase-1 levels were elevated in the HN878-infected macrophages, although each of these markers was differently expressed between these two groups at different time points post-infection (Figure 5A). This observation suggests intricate complexities in the regulation of NLRP3 activation in macrophages upon infection with different Mtb strains. We also observed a significantly higher level of caspase-activity in Mtb-infected rBMDMs, compared to the uninfected cells (Figure 5B). Importantly, the caspase activity was significantly higher in HN878-infected rBMDMs, compared to CDC1551-infected cells at 24 hpi. Together, these observations suggest upregulation of HIF*1A* expressions and inflammasome activation pathway during progressive Mtb infection in vivo (lungs) and in vitro (rBMDM and hu-MΦ).

**Figure 5.**
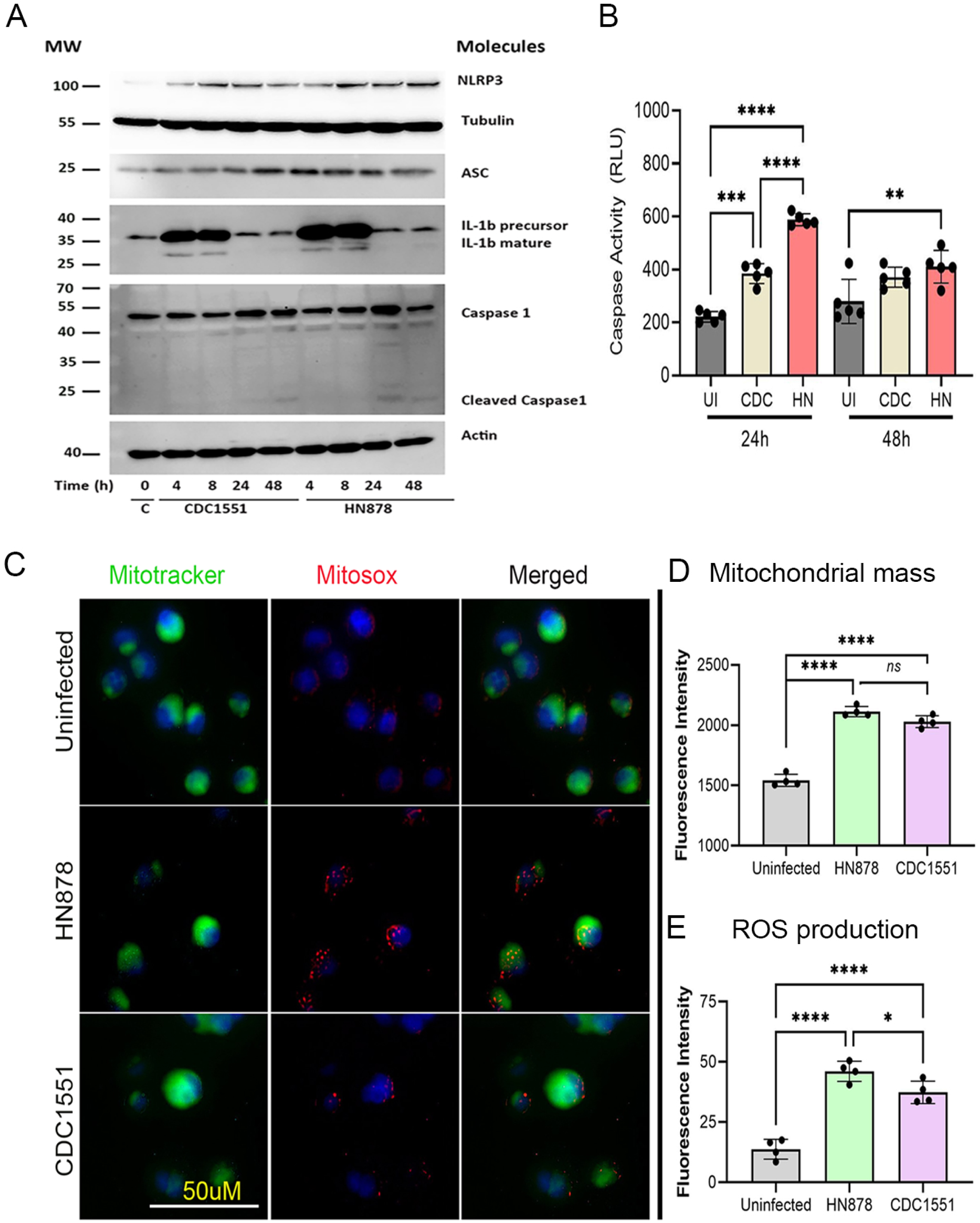
Production of NLRP3 inflammasome pathway proteins and differential activation of mitochondrial stress in macrophages during Mtb HN878 or CDC1551 infection. (**A**) Representative Western blot image of proteins isolated from HN878- or CDC1551-infected THP-1 showing levels of NLRP3, ASC, IL1β and Caspase 1 at 4, 8, 24 and 48 hpi. Uninfected cells (0, C) were used as controls. Tubulin and Actin were used to normalize samples. The IL1β panel shows both precursor and mature forms of the protein. The caspase 1 panel shows both uncleaved and cleaved forms of the protein. MW-molecular weight standard in kDa. (**B**) Fluorescence measurement of caspase activity in rBMDMs infected with CDC1551 or HN878 for 24h or 48h. (**C**) Representative images of uninfected or Mtb-infected THP-1 macrophages after staining with MitoTracker (green) and Mitosox (red). The scale bar is 50µm. (**D**) Quantifications of mitochondrial mass by MitoTracker staining (**D**) Quantifications of mitochondrial ROS production by MitoSOX staining. Fluorescence intensities of MitoTracker (mitochondrial mass) and MitoSOX (mitochondrial ROS) in hu-MΦ infected with HN878 or CDC1551 or no infection (Uninfected) were measured. Experiments were repeated twice with n=3-5 samples and statistical analyses were performed using one-way ANOVA. *P<0.05; ** P< 0.01; ***P<0.005; ****P<0.001.

### 2.6. Different mitochondrial stress responses in macrophages infected with Mtb HN878 and CDC1551

NLRP3 inflammasome activation has been shown to be triggered by mitochondrial stress (Liu et al., 2018). Therefore, we investigated the impact of Mtb infection on mitochondrial stress and associated activation of NLRP3 inflammasome pathway. Using MitoTracker Green and MitoSOX Red probes we determined mitochondrial mass and mitochondrial reactive oxygen species (ROS) production, respectively, in hu-MΦ with or without Mtb infection (Figure 5C). We found a significantly increased mitochondrial mass and ROS production in Mtb-infected hu-MΦ, compared to uninfected controls (Figure 5D and E). Importantly, HN878-infected hu-MΦ had a higher mitochondrial mass and ROS levels than CDC1551-infected cells, although only the difference in ROS levels was statistically significant between these two groups (Figure 5D and E). These observations suggest that progressive Mtb infection is associated with elevated mitochondrial dysfunction, particularly through ROS production in hu-MΦ, which can contribute to NLRP3 inflammasome activation.

### 2.7. Differential macrophage death upon infection by Mtb HN878 and CDC1551

The nature of infecting Mtb strain differentially induces the death of infected phagocytes by multiple mechanisms, leading to different extents of host inflammation and tissue destruction (Nisa et al., 2022). Since we observed differential regulation of inflammatory response and inflammasome activation network between HN878- and CDC1551-infected macrophages, we examined whether this would result in different host cell deaths. We measured the extent of apoptosis and necrosis in rBMDMs and THP-1 macrophages with or without Mtb infection by a fluorescence-based cellular assay (Figure 6). As expected, significantly higher percentages of apoptosis and necrosis were noted in Mtb-infected rBMDMs and THP-1 macrophages, compared to the uninfected controls (Figures 6A-D). The extent of both apoptosis and necrosis were proportional to the bacterial multiplicity of infection (MOI) used to infect macrophages. Importantly, at an MOI of 1 and 5, a significantly increased apoptosis and necrosis were noted in HN878-infected macrophages, compared to CDC1551-infected cells (Figures 6A-D).

**Figure 6.**
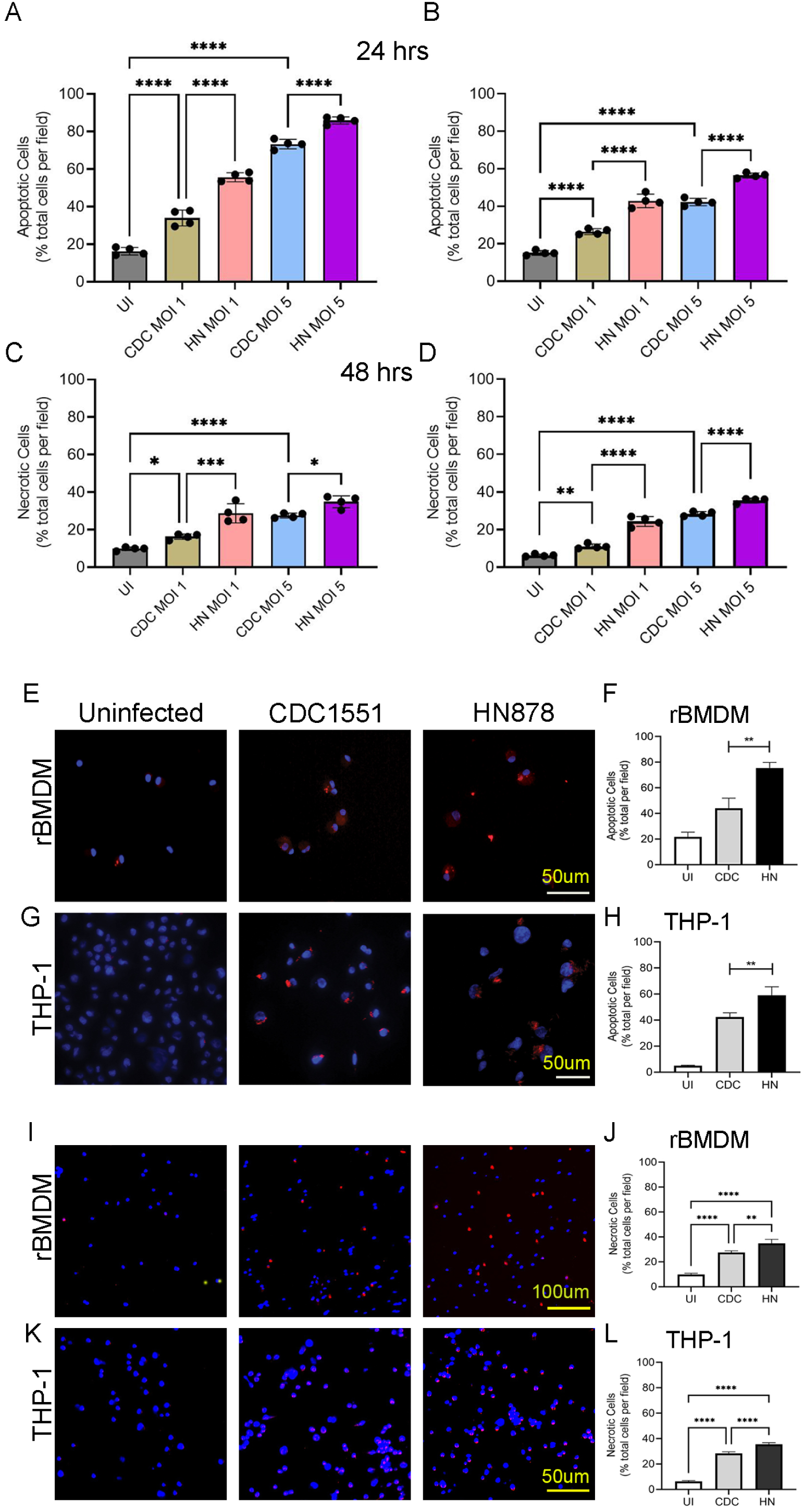
Apoptosis and necrosis of Mtb HN878- or CDC1551-infected macrophages. (A-. **D)** Percentages of apoptotic (**A, B**) or necrotic (**C, D**) rBMDMs (**A**, **C**) and THP-1 macrophages (**B**, **D**) infected with CDC1551 or HN878 at an MOI of 1 or 5 were determined at 24 h or 48h post infection. Uninfected (UI) cells were included as controls. (**E-L**) Representative images (**E**, **G**, **I**, **K**) and quantifications (**F**, **H**, **J**, **L**) of apoptotic (**E-H**) and necrotic (**I-L**) rBMDMs (**E**, **F**, **I**, **J**) and hu-MΦ (**G**, **H**, **K**, **L**) using immunocytochemistry. Cells were enumerated manually from the enlarged microscopic images of each field. Experiments were repeated thrice with n=3 samples, and statistical analyses were performed using one-way ANOVA. *P<0.05; ** P< 0.01; ***P<0.005; ****P<0.001.

To further validate and confirm differences in macrophage apoptosis and necrosis during HN878 and CDC1551 infections, we performed immunocytochemistry analysis using rBMDMs and THP-1 macrophages with or without Mtb infection. Consistent with the fluorescence-based cellular assay, the imaging analysis also showed significantly elevated apoptosis (Figures 6E-H) and necrosis (Figures 6I-L) of Mtb-infected macrophages, compared to untreated control cells. Importantly, apoptosis and necrosis were significantly higher in HN878-infected macrophages, compared to CDC1551 infection. Together, these observations indicate that progressive infection caused by HN878 is characterized by elevated death of infected macrophages.

### 2.8. Differential expression of GBP family genes between progressive and non-progressive Mtb infection in rabbit lungs

GBPs are among the top family of genes induced by IFNγ, which is crucial for Mtb control in vivo (Tretina et al., 2019; Olive et al., 2023; Diatlova et al., 2023; Ghanavi et al., 2021; Kim et al., 2011). The transcriptome of rabbit lungs infected with HN878 or CDC1551 for 4 weeks showed striking differences in the expression of member genes of the IFN-γ signaling (Supplementary Figure 5). The list of SDEGs in this pathway includes *IFNG*, *GBP1*, *IFIT1*, *IFIT1B*, *IRF1*, *IRF5*, *IRF7*, and *IRF8* as well as *STAT1*, encoding the transcriptional regulator of IFN signaling STAT 1 (Supplementary Figure 5A and B). Importantly, the expression of all genes in the IFN signaling pathway, except for *IRF6*, was upregulated in HN878-infected rabbit lungs (Supplementary Figure 5B) whereas most of these genes were either downregulated or not expressed to significant levels in the CDC1551-infected rabbit lungs (Supplementary Figure 5B). Importantly, *GBP1* was among the top upregulated SDEGs in HN878-infected rabbit lungs. Because GBPs play an important role in TB pathogenesis, we first profiled the expression of GBPs 1-5 in HN878 or CDC1551 infected rabbit lungs, rBMDM, and THP-1 macrophages by qPCR analysis. We observed a significantly upregulated expression of *GBP1* and *GBP2* in HN878-infected, compared to CDC1551-infected, rabbit lungs, rBMDM and hu-MΦ (Figure 7). Among the other GBPs tested, expression levels of *GBP4* and *GBP5* were significantly upregulated in rBMDM (at 48 hpi) and THP-1 cells (at 24 hpi) during HN878 infection, compared to CDC1551 infection, and the expression pattern of *GBP3* was inconsistent between Mtb-infected macrophages and infected rabbit lungs (Figure 7). A similar differential expression pattern of tested GBP genes was noted in between CDC1551 and H37Rv infected macrophages (Supplementary Figure 6). Taken together, among various GBPs tested, we observed *GBP1* to be consistently and significantly differentially expressed between HN878 and CDC1551 or H37Rv infections in the lungs, rBMDMs and in THP-1 macrophages.

**Figure 7.**
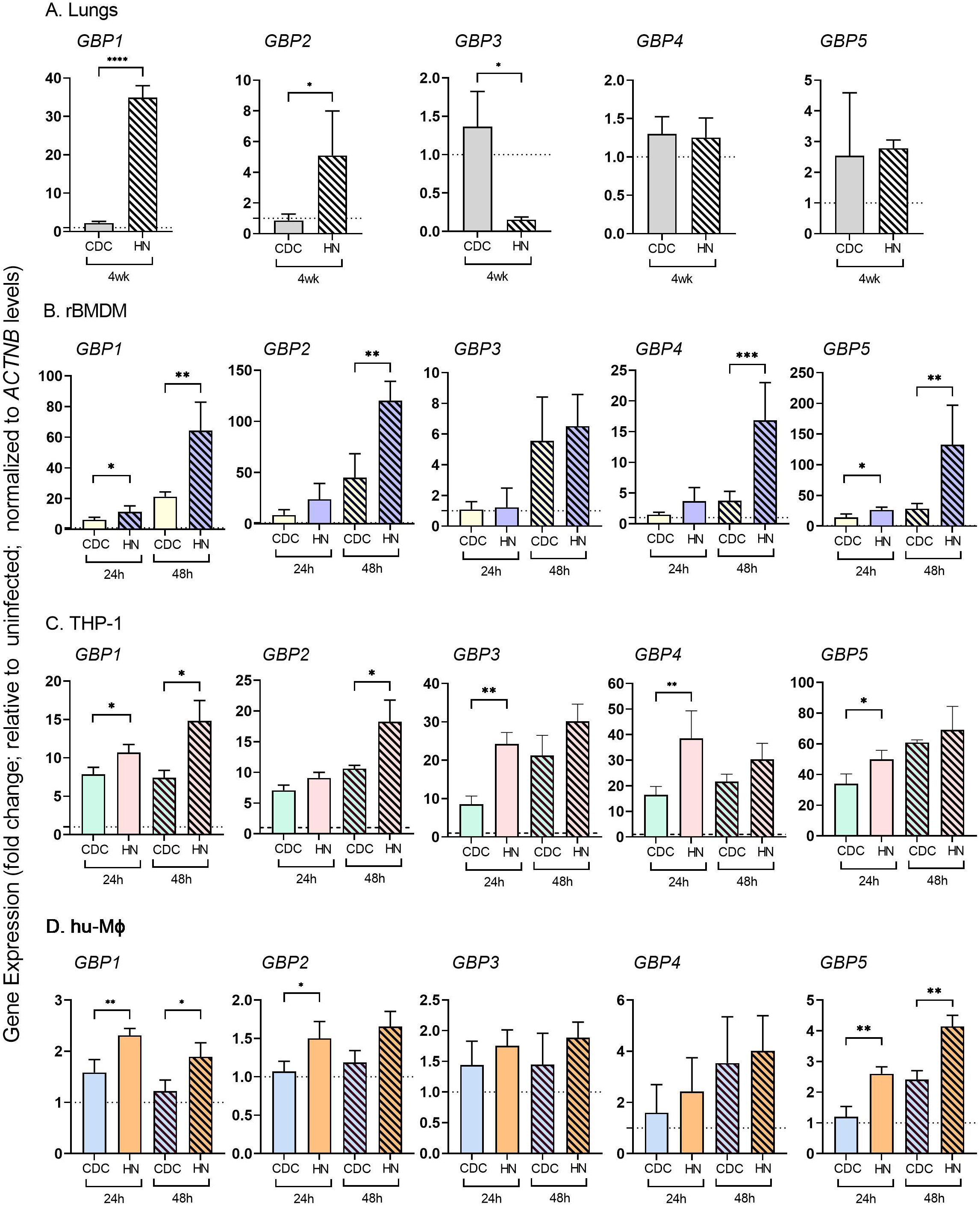
Expression profile of GBP family genes in macrophages during Mtb HN878 and CDC1551 infection. Transcript levels of *GBP1*, *GBP2*, *GBP3*, *GBP4* and *GBP5* in rabbit lungs 4 weeks post-infection (**A**) or in rBMDMs (**B**) or THP-1 (**C**) or hu-MΦ (**D**) infected with CDC1551 (CDC) or HN878 (HN) for 24 or 48 hours measured by qPCR. Fold changes of gene expression levels in Mtb-infected samples relative to uninfected samples are calculated. The level of *ACTNB* expression is used to normalize the expression level of test genes and the results are expressed as relative fold change. Experiments were repeated thrice with n=3-4 samples and statistical analyses were performed using the student *t*-test. *P<0.05; ** P < 0.01; ***P<0.005; ****P<0.001.

### 2.9. Differential regulation of inflammasome activation pathway by HIF1α and GBP1 in macrophages infected with different Mtb strains

GBP1 and the HIF-1α signaling pathways are involved in inflammasome activation in macrophages (Meunier et al., 2014; Pilla et al., 2014) (Jiang et al., 2020; Ouyang et al., 2013). We observed differential expression of NLRP3 inflammasome activation, GBP1 and HIF-1α pathway genes between HN878- and CDC1551-infected lungs and macrophages. Based on these findings, we hypothesized that GBP1 and HIF1α differentially regulates NLRP3 inflammasome activation between HN878- and CDC1551-infected macrophages. To test this hypothesis, we created THP-1 macrophages that are knocked down (KD) for *HIF1A* or *GBP1* expression using respective siRNAs and measured the expression of NLRP3 inflammasome activation markers and GBP family genes after infecting the cells with HN878, CDC1551 or H37Rv (Figure 8 and Supplementary Figure-7).

**Figure 8.**
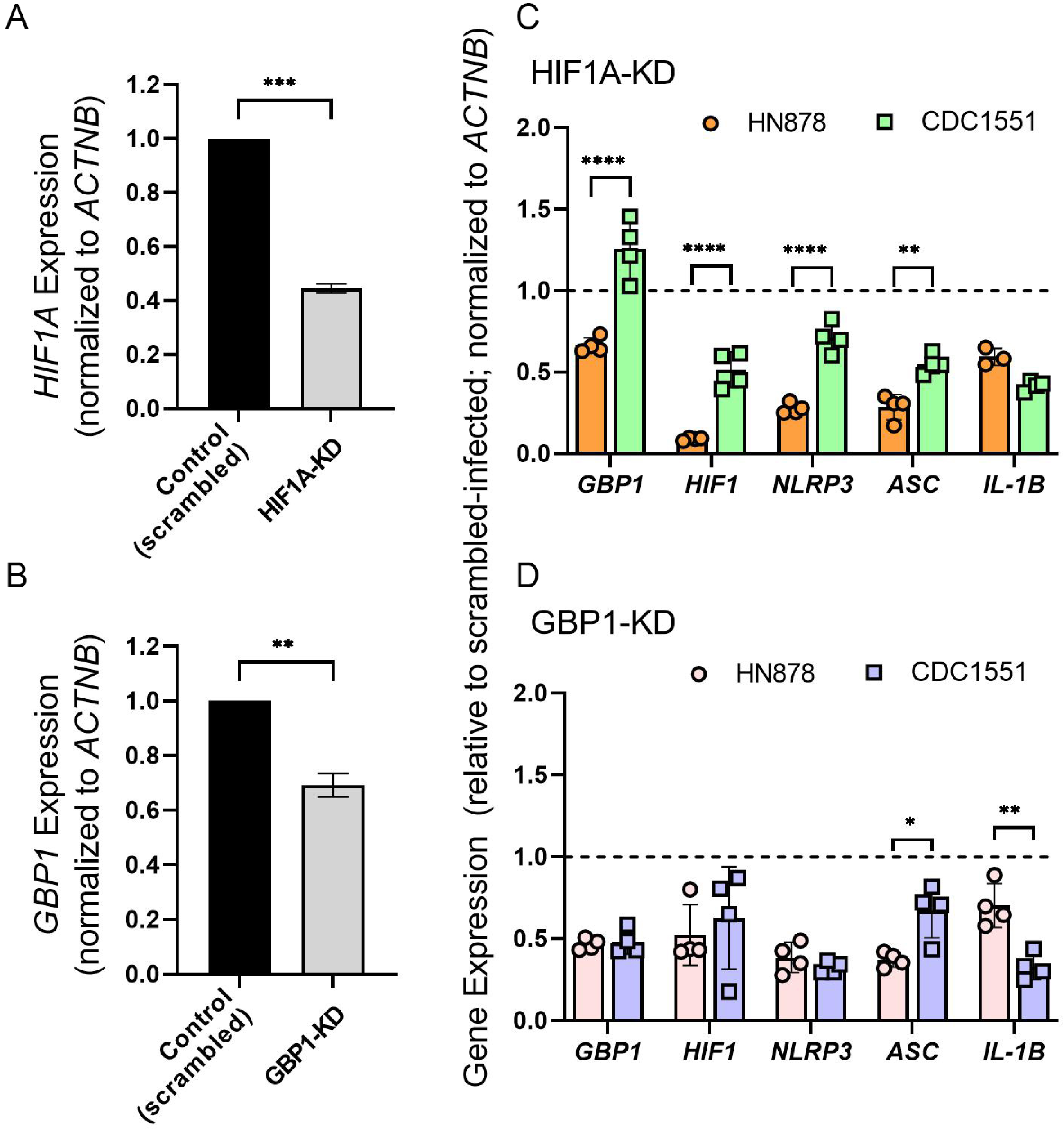
Regulation of NLRP3 inflammasome complex by HIF1A and GBP1 in macrophages during Mtb HN878 and CDC1551 infection. THP-1macrophages were treated with siRNAs against *HIF1A* (HIF1-KD) or *GBP1* (GBP1-KD) or control (scrambled) siRNA and the expression level of *HIF1A* (**A**) or *GBP1* (**B**) in these cells was determined. Macrophages treated with siRNAs against *HIF1A* or *GBP1* were infected with CDC1551 or HN878 and harvested at 24 h post-infection (**C** and **D**). Expressions of indicated marker genes (*GBP1*, *HIF1A*, *NLRP3*, *ASC*, *IL1B*) in harvested cells are measured by qRT-PCR, cells transfected with scrambled siRNA and infected with same Mtb strain were used as controls, the expression level of each gene in the control cells is set to 1 (dotted lines in **C** and **D**) and the expression level in siRNAs transfected and Mtb-infected samples is expressed as relative fold change. Expression level of *ACTNB* in each sample was used to normalize the expression of test genes in respective samples. Experiments were repeated thrice with n=3-4 samples and statistical analyses were performed using the student *t*-test. *P<0.05; ** P < 0.01; ***P<0.005; ****P<0.001.

The HIF1-KD and GBP1-KD macrophages had about 60% and 35% reduced expression of respective genes, compared to the control cells transfected with scrambled siRNA (Figures 8A, B). Mtb infection of both HIF1-KD and GBP1-KD macrophages dampened the expression of *GBP1*, *HIF1A*, *NLRP3, ASC* and *IL1B* in both GBP1-KD and HIF1A-KD macrophages, compared to control (scrambled siRNA) treated Mtb-infected macrophages (Figure 8C-D). A significant downregulation of *HIF1A*, *NLRP3* and *ASC*, and upregulation of *GBP1* was observed in HIF1A-KD macrophages, while downregulation of *ASC* and *IL1B* was noted in GBP1-KD macrophages infected with HN878, compared to CDC1551 infection (Figure 8C, D). Similarly, expression of *GBP1*, *NLRP3* and *IL1B* was downregulated in both HIF1-KD and GBP1-KD macrophages, while expression of *ASC1* and *HIF1A* was upregulated in HIF1-KD and GBP1-KD macrophages, respectively upon H37Rv infection, compared to controls (Supplementary Figure-7). The qPCR results were further confirmed by imaging analysis of respective macrophages (Figure-9A-E). In the control macrophages that were treated with scrambled siRNA the percentage of NLRP3 and IL1β positive cells were significantly increased upon Mtb infection (Figure 9A, D, E). Among these macrophages, the percentage of NLRP3 and IL1β positive cells were significantly higher during HN878-infection, compared to CDC1551-infection (Figure 9A, D, E). However, in both HIF1-KD and GBP1-KD macrophages, the percentage of NLRP3 and IL1β positive cells was significantly dampened following Mtb infection, compared to the uninfected controls (Figure 9B-E). There was no significant difference in the percentage of NLRP*3* and IL1β positive cells between HN878- and CDC1551-infected HIF1-KD and GBP1-KD macrophages or among uninfected control, HIF1-KD and GBP1-KD macrophages (Figure 9D, E). Together, these data indicate that both HIF1α and GBP1 are involved in inflammasome activation in THP-1 macrophages upon infection, although it may be independent of the nature of infecting Mtb strain.

**Figure 9.**
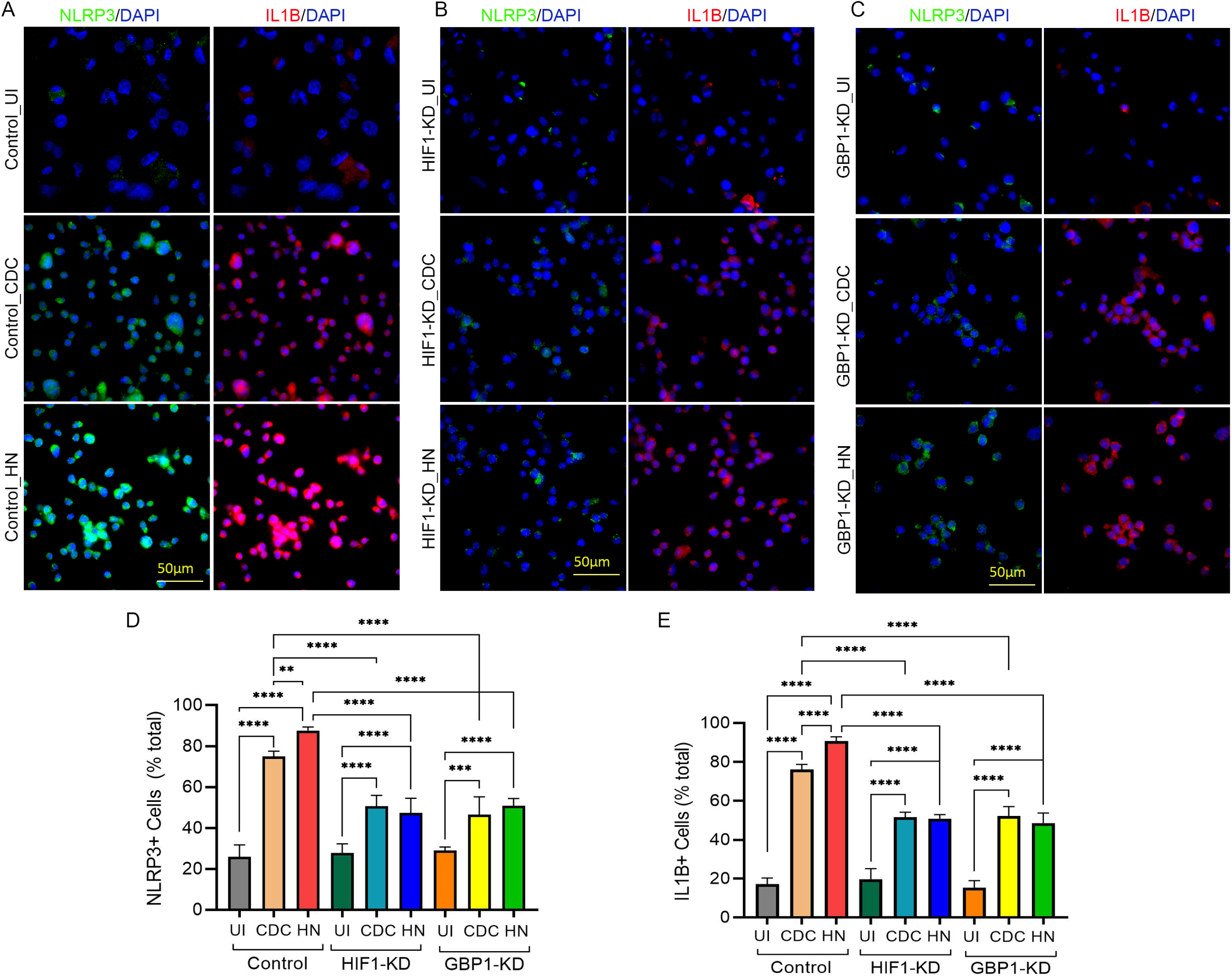
Regulation of expressions of Inflammasome markers by HIF1A and GBP1 in Hu-MΦ infected with Mtb HN878 or CDC1551. (**A-C**) THP-1 macrophages were treated with scrambled siRNA (Control, A) or siRNAs against *HIF1A* (B) or *GBP1* (C). Transfected cells were infected with Mtb and immunocytochemistry was performed using an antibody against NLRP3 or IL1β as described in the method section. (D and E) Cells expressing NLRP3 (**D**) or IL1β (**E**) were enumerated. Data are representative of experiments repeated twice with n=3-4 samples and statistical analyses were performed using one-way ANOVA. *P<0.05; ** P< 0.01; ***P<0.005; ****P<0.001.

Next, we determined the protein levels of pro- and anti-inflammatory cytokines and chemokines in control, HIF1-KD and GBP1-KD THP-1 macrophages following Mtb infection (Figure 10). In control macrophages, both HN878 and CDC1551 infections significantly increased the levels of proinflammatory (TNFα, IL1β, IL6, CXCL8) and anti-inflammatory (IL4, IL10) molecules and GMCSF. However, IL2 was significantly induced only in HN878-infected cells, compared to the uninfected controls. Compared to Mtb-infected control macrophages, the level of IL1β was significantly reduced in both HIF1-KD and GBP1-KD cells after infection with HN878 or CDC1551, while TNFα and IL6 were significantly reduced only in HN878-infected HIF1-KD and GBP1-KD macrophages. Similarly, GMCSF level was significantly reduced in HN878-infected HIF1-KD, compared to the infected control macrophages (Figure 10). However, no significant difference was observed in the levels of CXCL8, IL2, IL4 and IL10 between HN878- and CDC1551-infected HIF1-KD and GBP1-KD macrophages. Thus, the levels of various pro- and anti-inflammatory cytokines and chemokines are differently impacted in HIF1-KD and GBP1-KD macrophages after infection with HN878 or CDC1551. These observations also highlight the strong association between proinflammatory cytokine (IL1β, TNFα and IL6) production and HIF-1 and GBP1, particularly during HN878 infection.

**Figure 10.**
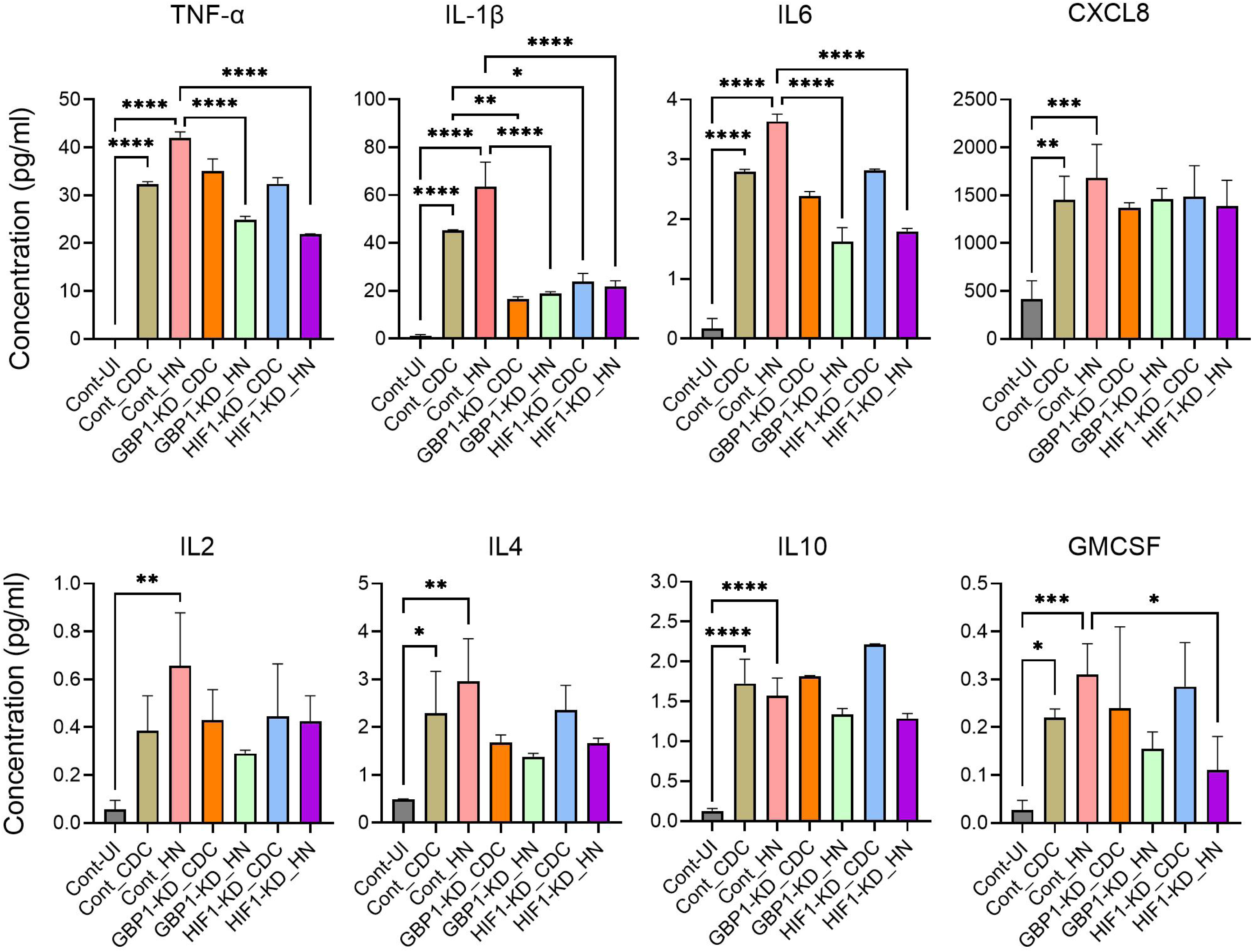
GBP1 and HIF-1A differentially regulate inflammatory cytokine production in Mtb HN878- or CDC1551-infected macrophages. THP-1 macrophages were treated with specific siRNAs against *GBP1* or *HIF1A* or with scrambled siRNA (Control) then infected with CDC1551 or HN878, and the cell-free supernatant was harvested at 48 hpi. Uninfected cells (UI) and cells treated with scrambled, non-specific siRNA and infected with Mtb strains (Cont) were used as controls. Luminex multiplex ELISA was used to determine the levels of TNF, IL1β, IL6, CXCL8, IL2, IL4. IL10, and GMCSF in the test and control samples. Experiments were repeated twice with n=3-4 samples and statistical analyses were performed using one-way ANOVA. *P<0.05; ** P< 0.01; ***P<0.005; ****P<0.001.

### 2.10. HIF1α and GBP1 differentially regulate macrophage death modalities during Mtb infection

To evaluate whether HIF1α and/or GBP1 were involved in the differential regulation of autophagy, apoptosis and necrosis between HN878- and CDC1551-infected macrophages, we infected HIF1-KD, GBP1-KD, and control THP-1 macrophages with HN878 or CDC1551 and enumerated the extent of autophagy, necrosis and apoptosis (Figure 11). We observed a significant increase in autophagy, necrosis and apoptosis in Mtb-infected control macrophages treated with scrambled siRNA, compared to the uninfected macrophages (Figure 11A-C). Compared to the Mtb-infected control macrophages, KD of either GBP1 or HIF1 completely abolished the induction of necrosis by both Mtb strains, reduced the induction of apoptosis by both Mtb strains, and reduced LC3 expression, a marker of autophagy, induced by HN878 (Figure 11A-C and Supplementary Figure 8). In addition, KD of GBP1, but not KD of HIF1, reduced autophagy induced by CDC1551 infection. Only KD of GBP1 differentially affected the induction of autophagy between CDC1551 and HN878 infection marked by LC3 expression (Figure 11A and Supplementary Figure 8). Thus, HIF1α and GBP1 differently regulate autophagy, necrosis and apoptosis during Mtb infection of THP-1 macrophages, although the infecting Mtb strain does not significantly impact the outcome of HIF1α and/or GBP1-mediated cell death mechanisms.

**Figure 11.**
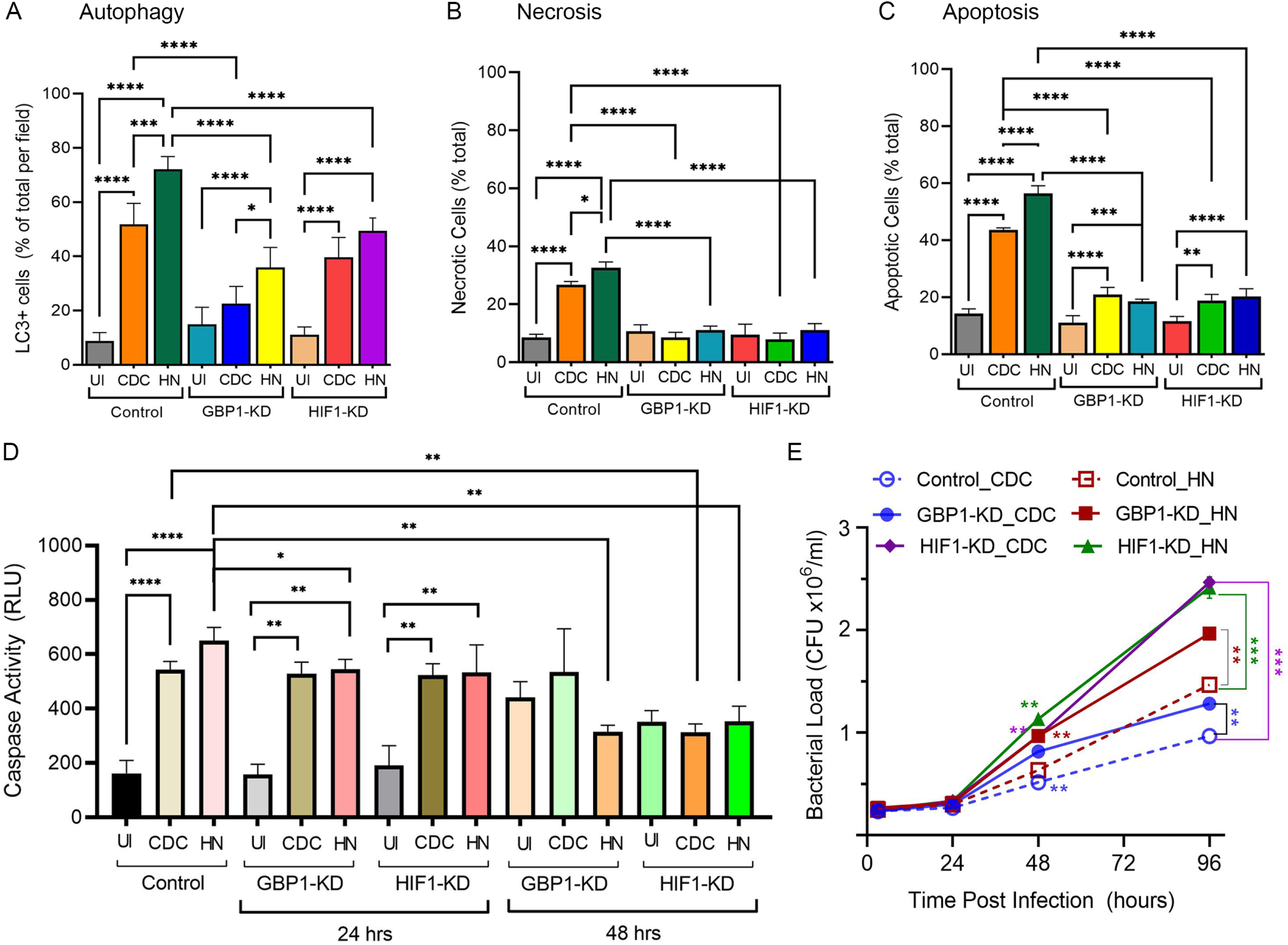
GBP1 and HIF-1A differentially regulate Caspase 1 activity and host cell death and mycobacterial proliferation in Mtb HN878 -and CDC1551-infected macrophages. (A-. **D)** THP-1 macrophages were treated with specific siRNAs against *GBP1* or *HIF1A* or with scrambled siRNA (Control) and then infected with CDC1551 or HN878. The percentage of autophagic (**A**), necrotic (**B**), and apoptotic (**C**) cells were enumerated by fluorescence-based microscopic imaging analysis and calculated as the percentage of total cells in the field. and the caspase activity in these cells was enzymatically measured (**D**). (**E**). The proliferation of Mtb strains in THP-1 macrophages treated with siRNAs against *GBP1* or *HIF1A* and then infected with CDC1551 or HN878 was determined. Uninfected cells (UI) and cells treated with scrambled, non-specific siRNA and infected with Mtb (Control) were used as controls. Experiments were repeated twice with n=3-4 samples and statistical analyses were performed using one-way ANOVA. *P<0.05; ** P< 0.01; ***P<0.005; ****P<0.001.

Next, we compared the caspase activity in HIF1-KD and GBP1-KD after HN878 or CDC1551 infection, to those in the uninfected, as well as infected control macrophages (Figure 11D). Similar to our observations with rBMDMs, Mtb-infection significantly increased the caspase activity in control macrophages (scrambled siRNA treatment), compared to the uninfected cells at 24 hpi. Similarly, the caspase activity was significantly increased in Mtb-infected HIF1-KD and GBP1-KD, compared to uninfected KD cells, independent of the Mtb strain, particularly at 24 hpi (Figure 11D). However, the level of caspase activity in HN878-infected GBP1-KD hu-MΦ was significantly lower than that in the infected control cells (scrambled siRNA treated). At 48 hpi, no significant differences were noted in the caspase activities between the uninfected and Mtb-infected GBP1- or HIF1-KD macrophages. However, at this time point, both CDC1551- and HN878-infected HIF1-KD showed significantly reduced caspase-1 activities, compared to the infected control cells (scrambled siRNA treated) (Figure 11D). Together, these observations suggest that HIF1A and GBP1 regulate the caspase-1 activity differently between HN878- and CDC1551-infected THP-1 macrophages, which is consistent with the results of apoptosis assay.

Finally, we tested whether HIF1α and/or GBP1 differentially affects the intracellular survival of HN878 and CDC1551 in macrophages. As shown in Figure 11E, a significantly higher bacillary load was noted in control (scrambled-RNA treated) macrophages infected with HN878, compared to CDC1551, starting at 48 hpi, until 96 hpi. Compared to the control, HIF1-KD and GBP1-KD macrophages had a significantly higher bacterial load upon infection with HN878 or CDC1551 at 48h and 96h post-infection (Figure 11E). Although the bacterial load was higher in HN878-infected HIF1-KD and GBP1-KD, compared to CDC1551-infection, the difference was significant only in GBP-KD at 96 hpi (Figure 11E). Together, these observations suggest that HIF1α and GBP1 differentially contribute to controlling intracellular Mtb growth in THP-1 macrophages.

## 3. DISCUSSION

Clinical studies have demonstrated the presence of heterogeneous Mtb strains with variable virulence and transmission potentials in the sputum of patients with active pulmonary TB (Dannenberg and Collins, 2001; Dusthackeer et al., 2019; Hunter, 2020; Verma et al., 2019). Because the nature of Mtb strains impacts the overall immunity and response to anti-TB therapy in patients, it is important to understand the mechanism by which various clinical Mtb strains impart variable host responses (Dannenberg and Collins, 2001; Dusthackeer et al., 2019; Hunter, 2020; Verma et al., 2019). The outcomes of Mtb infections in model animals of TB is affected by the nature of infecting Mtb strain, ranging from LTBI to active TB (European Concerted Action on New Generation Genetic et al., 2006; Koo et al., 2012; Subbian et al., 2013). At the molecular level, the pathogen-associated molecular patterns of these Mtb strains and their interactions with pattern recognition receptors (PRRs) of host cells regulate subsequent cellular signaling, leading to differences in the activation of various host cell processes (European Concerted Action on New Generation Genetic et al., 2006; Liu et al., 2017). These biological processes, including apoptosis, necrosis and autophagy, differentially contribute to the progressive or non-progressive Mtb infection (European Concerted Action on New Generation Genetic et al., 2006; Liu et al., 2017).

In this study, taking cues from genome-wide transcriptome data of human and rabbit lungs, we report that macrophage inflammasome activation is differentially regulated by HIF-1α and GBP1 between progressive and non-progressive Mtb infections. These findings are consistent with and supported by *in vitro* studies conducted using two clinical Mtb strains that are known to elicit different immune responses in macrophages and animal models (Koo et al., 2012; Manabe et al., 2003; Subbian et al., 2012; Subbian et al., 2011; Tsenova et al., 2020). We observed a markedly higher activation of the inflammasome in response to infection with the progressive disease-causing *Mtb* strain HN878, compared to the non-progressive strain CDC1551. This differential inflammasome activation was accompanied by a robust induction of several key effector pathways, including apoptosis, necrosis, and autophagy, particularly in the host tissues harboring HN878. These findings suggest that the pathogenicity of *Mtb* strains may be closely tied to their ability to manipulate host cell death pathways, with more virulent strains, such as HN878, triggering a more intense immune response that accelerates disease progression. In the case of HN878, the higher levels of inflammasome activation may promote a more intense inflammatory response, which could drive the formation of necrotic granulomas and the breakdown of lung tissue, facilitating the progression from latent to active TB. This is in stark contrast to the response elicited by CDC1551, where the inflammasome response was more controlled, likely contributing to the establishment of latent infection and a more contained disease state.

The inflammasome complexes play a key role in regulating the immune response, including inflammation, during infections (Barnett et al., 2023; Huang et al., 2019; Man et al., 2016; Rastogi and Briken, 2022). Recognition of Mtb by PRRs during phagocytosis activates NLRP3 inflammasome complex, which helps in the fusion of Mtb-containing phagosomes with lysosomes, and control of bacterial growth in macrophages (Barnett et al., 2023; Chen and Ichinohe, 2015; da Costa et al., 2019; Jayaraman et al., 2013; Ma et al., 2021; Pilli et al., 2012; Rastogi and Briken, 2022; Subbarao et al., 2020; Yen et al., 2015). Particularly, the phenolic glycolipids of HN878 were shown to activate the inflammasome in murine phagocytes (Domingo-Gonzalez et al., 2017). On the contrary, clinical Mtb strains associated with severe disease in humans have been shown to evade killing by NLRP3 inflammasome activation-mediated effector functions in murine macrophages (Sousa et al., 2020; Subbarao et al., 2020). Thus, the signaling mechanisms responsible for the differential inflammasome activation in phagocytes upon infection with various clinical Mtb strains are complex and are not fully understood. One of the plausible explanations for the differential regulation of inflammasome activation involves the expression of IL1β, which is associated with host protective and permissive responses during Mtb infection (Rastogi and Briken, 2022). Thus, a complex interaction of inflammasome activator and effector molecules defines cellular fate. In this study, we report HIF-1α and GBP-1 as two of the factors differentially regulate NLRP3 inflammasome activation in macrophages during Mtb infection, perhaps through IL1β mediated mechanism.HIF-1α is a key transcriptional regulator activated in response to hypoxia in cells and granulomas (Cardoso et al., 2015; Kumar et al., 2024; Resende et al., 2020). The role of HIF-1α in restricting mycobacterial growth is well documented (Li et al., 2021; McGettrick and O’Neill, 2020; Zenk et al., 2021). In a recent study, it was found that the stabilization of HIF-1α in human macrophages led to reduced Mtb growth through the activation of Vitamin D-mediated antimicrobial pathway (Zenk et al., 2021). Similarly, HIF-1α contributes to the IFN-γ-dependent control of Mtb infection in murine macrophages and mouse lungs (Braverman et al., 2016). Consistently, mice deficient in HIF-1α were more susceptible to mycobacterial infection than wild-type animals (Braverman et al., 2016; Cardoso et al., 2015). The HIF-1α expression was also reported to track the disease progression in a BALC/c murine model of Mtb infection; HIF-1α level was also induced in activated macrophages during the early stages of Mtb infection but the level was much higher in foamy macrophages during chronic stages of infection (Baay-Guzman et al., 2018). Furthermore, the blockade of HIF-1α impaired the inflammasome activation and proinflammatory responses of Mtb-infected hu-MΦ (Braverman et al., 2016; Cardoso et al., 2015). Taken together, these observations suggest that HIF-1α regulates TB pathogenesis by several factors, including the host cell activation status at the time of infection and the nature of infecting Mtb strain. Consistent with these studies, we found increased HIF-1α expression concomitantly with elevated NLRP3 inflammasome activation network in rabbit and human lungs with necrotic granulomas and inflammation. Results presented here further suggest that the infecting Mtb strain differentially affects the expression of HIF-1α and NLRP3 inflammasome activation network genes and contributes to differential outcomes of infection (i.e., progressive vs. non-progressive).

We observed that GBPs play a key role in differentially regulating inflammasome activation in macrophages during progressive or non-progressive Mtb infection. GBPs are known to play a key role in regulating host innate immune response during bacterial infections via inflammasome activation, induction of oxidative responses, and autophagy (Costa Franco et al., 2018; Cui et al., 2021; Fisch et al., 2019; Gomes et al., 2019; Kim et al., 2016; Kim et al., 2012; Kim et al., 2011; Liu et al., 2018; Marinho et al., 2020; Olive et al., 2023; Qiu et al., 2018; Santos et al., 2020; Shenoy et al., 2012; Tretina et al., 2019). Specifically, GBP1 was highly expressed in the blood cells of TB patients and its expression pattern correlated with that of other IFN-stimulated genes (Shi et al., 2022). GBP1 was also reported to be differentially expressed between active TB and LTBI cases, suggesting the potential of this molecule as a diagnostic biomarker (Peer et al., 2023). Consistent with these reports, we observed upregulated GBP1 expression in rabbit and human lungs with progressive TB and HN878-infected primary macrophages. Furthermore, using the siRNA-knockdown approach, we demonstrated that GBP1 differentially regulates inflammasome activation and subsequent proinflammatory molecule production in macrophages infected with HN878 or CDC1551.

Finally, we observed that progressive and non-progressive infections induce different host cell death pathways in rabbit and human lungs. Higher percentages of cells positive for apoptosis, necrosis and autophagy were noted in the lungs with progressive disease and in HN878-infected macrophages. These results are consistent with and supported by previously published findings (Amaral et al., 2024; Roca et al., 2019). During Mtb infection, the host cell death caused by apoptotic, autophagic, and necrotic pathways have different outcomes for the progression of disease (Bertheloot et al., 2021). In general, apoptosis of infected macrophages is believed to kill Mtb whereas necrosis of infected macrophages is believed to promote the release of viable Mtb, further promoting progressive infection (Lam et al., 2017). However, highly virulent Mtb strains, such as HN878, can divert the apoptotic pathway and promote necrosis to exacerbate disease pathology in phagocytes (Behar et al., 2010; Blomgran et al., 2012; Butler et al., 2012; Nisa et al., 2022). While the role of autophagy in targeting and restricting intracellular Mtb growth is well established, the importance of autophagy in resisting Mtb infection in animal models of infection has been controversial (Behar and Baehrecke, 2015; Golovkine et al., 2023; Gutierrez et al., 2004; Kimmey et al., 2015). For example, ATG-5-mediated autophagy induction is differentially regulated between macrophages and neutrophils in a murine model of TB and the loss of ATG-5 negatively affects Mtb control in neutrophils but not in macrophages (Kimmey et al., 2015). Similarly, Mtb has evolved with mechanisms to resist the induction of autophagosome formation and/or its fusion with the lysosome to evade killing by the phagocytes (Songane et al., 2012). These studies indicate the complex interplay between various host cell death pathways and warrant further investigation to understand their contribution to progressive and non-progressive Mtb infections.

In summary, we unraveled a novel mechanism of inflammasome activation mediated by GBP1 and HIF-1α, which orchestrates differential immune response in macrophages and in vivo upon infection with Mtb strains that cause a progressive or non-progressive infection. These findings give new insight into the immunomodulatory activities of clinical Mtb strains, which may provide useful information about the key molecules, such as GBP1, that can be targeted for developing host-directed therapeutics to treat TB and preventing reactivation of LTBI (Antimicrobial Resistance, 2022). Understanding the intricate dynamics between inflammasomes, apoptosis, necrosis, and autophagy will be critical for advancing our ability to control TB and mitigate its devastating global impact.

## 4. MATERIALS and METHODS

### 4.1. Mycobacterial culture

Clinical *Mycobacterium tuberculosis* strains CDC1551 and HN878 and laboratory strain H37Rv were grown in Middlebrook 7H9 (BD Biosciences, Franklin Lakes, NJ, USA) medium supplemented with 10% ADC, 0.5% glycerol and 0.05% Tween 80 at 37°C. The culture was grown to logarithmic phase (OD ̴ 0.6) and stored at −80°C until further use. All chemicals were purchased from Sigma (Sigma-Aldrich, St. Louis, MO, USA) unless mentioned otherwise.

### 4.2. Rabbit aerosol infection

Female New Zealand white rabbits (*Oryctolagus cuniculus*) of ∼2.5 kg body weight was purchased from Covance Inc (Covance Research Products, Denver, PA). Rabbits were randomly assigned into groups and exposed to aerosols containing HN878 or CDC1551, as described previously (Subbian et al., 2011; Subbian et al., 2012). Five rabbits (n=5) per group (uninfected, HN878-infected or CDC1551-infected) per time point (T=0 and 4 weeks) were used in this study (total number of animals=30). All animal studies were approved by the Rutgers University Institutional Animal Care and Use Committee (IACUC).

### 4.3. Isolation, culture, and infection of rabbit bone marrow derived macrophages (rBMDM)

Femur bones of rabbits were used for rBMDM isolation as described earlier (Cody et al., 2005). Briefly, bones were cut open from ends and flushed using DMEM and an 18-gauge needle. The bone marrow obtained was passed through a 70 μM cell strainer and washed using PBS by centrifugation. The pellet obtained was lysed using ACK lysis buffer (ThermoFisher Scientific, Waltham, MA, USA), washed using PBS, and resuspended in DMEM. The cells were seeded at a density of 3×10^6^ cells/ml in culture dishes containing DMEM supplemented with 10% FBS and 40 ng/ml MCSF (Peprotech, Cranbury, NJ, USA). The cells were left for differentiation for 6 days at 37°C in a 5% CO_2_ incubator. Differentiated bone marrow-derived macrophages (BMDMs) were infected with Mtb strains at a multiplicity of infection (MOI) 5 for 2h. Extracellular bacteria were removed by washing thrice with sterile PBS and the cells were further incubated at 37°C for required time intervals. For CFU determination, infected cells were lysed using 0.06% SDS and diluted in PBS-tween. Diluted cultures were spread on 7H10 agar plates (BD Biosciences, Franklin Lakes, NJ, USA) supplemented with 10% OADC and 0.5% glycerol. Bacterial colonies were counted after 3-4 weeks of incubation.

### 4.4. Hu-MΦ culture, gene silencing and Mtb infection

Human peripheral blood monocytes (PBMCs) were isolated from heparinized whole blood from healthy donors by density gradient centrifugation as described by Kumar et al. (2015). Briefly whole blood was layered onto Ficoll-Paque PLUS (Cytiva, Wilmington, DE, USA) and centrifuged to obtain the mononuclear cell layer. The mononuclear cell layer was collected, washed, seeded in culture flasks and allowed to adhere for 2 h. Lymphocytes were washed away, and the adhered cells were allowed to differentiate for 7-10 days in RPMI supplemented with 20mM HEPES, 80 µM L-glutamine, 100 U/mL penicillin, 100 µg/mL streptomycin (Gibco, USA), 10% FBS and 40 ng/ml MCSF (Peprotech, Cranbury, NJ, USA). The day before infection, cells were seeded in 12 well culture plates (5 x 10^5^ cells/well) in complete RPMI medium free of antibiotics. Human monocytic THP-1 cells (TIB-202) were purchased from American Type Culture Collection (ATCC; Manassas, VA, USA), and cultured in RPMI 1640 medium (HyClone™ Logan, Utah, USA) supplemented with 10% (v/v) heat-inactivated fetal bovine serum (Gibco, Billings, MT, USA), 2 mM L-glutamine (Gibco, Billings, MT, USA), 100 U/mL penicillin, 100 µg/mL streptomycin (Gibco, USA), 10 mM 4-(2-hydroxyethyl)-1 piperazine-ethane-sulfonic acid (HEPES; HyClone™ Logan, Utah, USA) and 0.05 mM 2-mercaptoethanol (Bio-Rad, Hercules, CA, USA). Cells were maintained in a humidified incubator at 37°C with 5% CO_2_ and sub-cultured every 2–3 days. To differentiate into macrophages, the THP-1 cells were seeded at a density of 1×10^5^ cells/well in 96-well tissue culture plates (Nunc™, Waltham, MA, USA) or 2×10^6^ cells/well in 6-well plates (Nunc™, Waltham, MA, USA) and treated with 50 ng/mL PMA (Sigma-Aldrich, St. Louis, MO, USA) for 24 h at 37°C in 5% CO_2_. Cells were washed twice with PBS and maintained in antibiotic-free RPMI 1640 medium for 24 h at 37°C in 5% CO_2_ and used in subsequent experiments. For silencing, siRNAs against HIF1α and GBP1 (Santa Cruz Biotechnology, Dallas, Texas., USA) were transfected into hu-MΦ (THP-1) using Lipofectamine LTX (ThermoFisher Scientific, Waltham, MA, USA) following manufacturer’s instructions. The concentration of siRNAs used was 50nM and cells were kept for 48h for gene silencing. Differentiated macrophages were infected with Mtb strains at a multiplicity of infection (MOI)1-5 for 2h. CFU was determined using the procedure described in above section.

### 4.5. Western blot analysis

Western blot to analyze the production of proteins was carried out as described previously (Kumar et al., 2016). Briefly, THP1 macrophage cells were washed in ice-cold PBS and lysed in lysis buffer (Cell Signaling Technology, Beverly, MA, USA) supplemented with a complete protease inhibitor ‘cocktail’ (Sigma-Aldrich, St. Louis, MO, USA) for 20 min on ice. Lysates were centrifuged at (14,000 x g) for 10 min to remove cell debris and supernatant was collected. Further, the supernatant was boiled in SDS denaturing Laemmli buffer for 10 min, separated on SDS-PAGE, transferred to PVDF membranes, which were blocked for 1 h at room temperature in 5% (w/v) nonfat dry milk (NFDM) in TBST (20mM Tris-HCl pH7.5, 150mM NaCl and 0.1% Tween 20) and incubated at 4°C overnight with antibody at 1:1000 dilution or beta-actin antibody (Cell Signaling Technology, Beverly, MA, USA) at 1:4000 dilution. The membrane was washed thrice for 5 min each in TBST and further incubated in HRP-conjugated secondary antibody (1:2000 dilution) in 5% NFDM (w/v) in TBST. The blot was developed using HRP substrate (Amersham ECL ^TM^ Prime Western Blotting Detection Reagents, GE Healthcare, Little Chalfont, Buckinghamshire, UK) as per the manufacturer’s instructions. Antibodies against NLRP3, ASC, Caspase-1, Cleaved Caspase-1, IL1β, Cleaved IL1β, Tubulin and Actin B were used as suggested by the manufacturer (Cell Signaling Technology, Beverly, MA, USA). Unprocessed, raw images of the Western blots are presented in Supplementary Figure-9.

### 4.6. Host cell RNA isolation and quantitative Real-time PCR (qRT-PCR)

Total RNA was isolated from tissue samples, human PBMC derived macrophages, THP1 and rBMDMs using Trizol (ThermoFisher Scientific, Waltham, MA, USA). Briefly, cells or tissue samples were lysed with Trizol reagent, and total RNA was isolated using the RNeasy mini kit (Qiagen, Germantown, MD, USA) following the manufacturer’s instructions. For quantitative gene expression analysis, cDNA was synthesized using Revertaid first-strand cDNA synthesis kit (ThermoFisher Scientific, Waltham, MA, USA). The cDNA was amplified with gene-specific primers using a Syber green-based qRT-PCR kit (ThermoFisher Scientific, Waltham, MA, USA). The expression level of target genes was normalized with the expression level of the housekeeping beta-actin gene (*ACTNB*). Fold change in gene expression was determined using the formula 2^-ΔΔCt^, where Ct is the threshold cycle. Relative expression of genes in infected samples was compared with uninfected ones for the time points.

### 4.7. Immunostaining on Tissue Section and macrophages

Tissue sections were fixed using neutral buffered formalin for 3 weeks followed by their dehydration through a series of graded ethanol baths and embedding into paraffin wax. After sectioning, 5μm thick tissue sections were placed onto glass slides. The (Formalin fixed paraffin embedded) FFPE tissue sections were deparaffinized by dipping in xylene, followed by rehydration through washing in graded ethanol. Then the tissue sections were kept in citrate buffer at 90°C for 40 min to retrieve antigen. Macrophages were fixed by treating with 4% paraformaldehyde for 20 min followed by washing twice with 1X PBS. The cells were kept in 70% ethanol until processed for immunostaining. For staining with antibodies, the slides or macrophages were incubated in blocking buffer containing 2% BSA in PBS for 1 h followed by incubation with primary antibody diluted in blocking buffer (1:500) overnight. The slides or macrophages were washed with PBS and a secondary antibody diluted in blocking buffer (1:1000) was added for 1h. The slides were washed with PBS to remove unbound antibodies, treated with True Black, and mounted. Primary antibodies were used against HIF1A (ThermoFisher Scientific, Waltham, MA, USA), NLRP3 (Cell Signaling Technology, Beverly, MA, USA), IL1B (Cell Signaling Technology, Beverly, MA, USA) and LC3 (Cell Signaling Technology, Beverly, MA, USA). The secondary antibodies used were from Abcam (Waltham, MA, USA).

### 4.8. Imaging and analysis

Images were acquired using an Axiovert 200M inverted fluorescence microscope (Zeiss, Oberkochen, Germany) using 20X objective or 63X oil-immersion objective and a Prime sCMOS camera (Photometrics, Tucson, AZ) controlled by Metamorph image acquisition software (Molecular Devices, San Jose, CA). To compare and quantify fluorescence signals arising from image acquisition, ImageJ (National Institutes of Health, Bethesda, MD, USA) software was used.

### 4.9. Caspase Assay

Macrophages (rBMDM and hu-MΦ) were analyzed for measurement of the activity of caspase-1 in cells, following the manufacturer’s (Caspase-Glo1 Inflammasome Assay, Promega, Madison, WI, USA) instructions. Briefly, cells were seeded in a white opaque 96-well plate and infected with CDC 1551 or HN878. After 24/48h of infection, Caspase-Glo reagent was added in the wells. Cells were incubated for an hour and luminescence was measured on Promega GloMax® Plate Reader (Promega, Madison, WI, USA).

### 4.10. Multiplex Cytokine Assay

Cytokines in the culture supernatants were analyzed using the Human Cytokine Magnetic 10-Plex Panel (ThermoFisher Scientific, Waltham, MA, USA) following the manufacturer’s instructions. Briefly, culture supernatants were collected and passed through a 0.45 µ nylon filter (VWR International, Radnor, PA, USA). Antibody beads from the kit were diluted to 1X and 25 µl was added in each well of a 96-well plate, followed by the addition of culture supernatants or standard. Then the plate was incubated at 4°C overnight. The wells were washed twice and 100µl of Biotinylated Detector Antibody was added and the plate was incubated at room temperature for an hour. After washing twice, 100 µl of 1X Streptavidin-RPE was added, and the plate was incubated for 30 min. Finally, the plate was washed twice and read in the Luminex System. The data were analyzed using ProcarataPlex Analysis software (ThermoFisher Scientific, Waltham, MA, USA).

### 4.11. Apoptosis and Necrosis Assay

Macrophages (rBMDM and hu-MΦ) were analyzed for apoptotic and necrotic cell death following the manufacturer’s instructions (Biotium, Fremont, CA, USA). Briefly, cells were seeded in 96 well plates and infected. After 24h of infection, cells were washed with 1X Binding Buffer followed by washing with PBS. Staining solution was added, and cells were incubated at room temperature for 15min. Cells were washed again with 1X binding buffer and observed under a microscope.

### 4.12. Mitochondrial oxidative stress assay

Generation of mitochondrial reactive oxygen species (ROS), especially superoxide, in hu-MΦ macrophages during mycobacterial infection was detected using stains MitoTracker (Cell Signaling Technology, Danvers, MA, USA) and MitoSox (Thermo Fisher Scientific, Waltham, MA, USA) following manufacturers’ instructions. Briefly, hu-MΦ (THP1) were prepared as discussed above. Cells were plated in a 24-well plate at a density of 1X10^5^ /well. Macrophages were infected at an MOI of 5 for 2h, followed by incubation for another 24h in a humidified incubator at 37°C with 5% CO_2._ Macrophages were stained with 400nM of MitoTracker and 2.5 µM of Mitosox for 30min in the CO_2_ incubator, washed with Phenol Red free RPMI supplemented with 10% FBS, and kept in the same medium for further analyses. Fluorescence intensity was measured. To confirm mitochondrial superoxide generation, macrophages were also examined under a fluorescence microscope.

### 4.13. Statistical analysis

All experiments were conducted in a minimum of three biological replicates, and the average value of two technical replicates was used for graphical presentations as the mean ± standard error values. Comparisons between two experimental conditions were analyzed by unpaired *t*-test with Welsh correction, and for multiple group comparison, one-way ANOVA with Tukey’s correction or two-way ANOVA was used. Statistical analysis was performed using GraphPad Prism 9.3 version (GraphPad Software, La Jolla, CA). For all the experimental data comparisons between groups, differences were considered statistically significant when *P* ≤ 0.05.

## Supporting information

Supplemental Files_Combined

## 5. SUPPLEMENTARY MATERIALS

**Supplementary Figure 1.** Proliferation of Mtb H37Rv, HN878 and CDC1551 in rBMDM and hu-MΦ and differential regulation of canonical immunological pathways in rabbit lungs infected with Mtb HN878 and CDC1551.

**Supplementary Figure 2.** Expression of genes involved in immune pathways in the lungs of patients with TB and caspase-activity in Mtb-infected rBMDMs

**Supplementary Figure 3.** Expressions of genes in the canonical inflammatory cytokine/chemokine signaling pathway in Mtb-infected rabbit lungs.

**Supplementary Figure 4.** Expression profile of NLRP3 inflammasome activation pathway genes in macrophages during Mtb H37Rv infection.

**Supplementary Figure 5.** Expression profile of IFN signaling pathway genes in rabbit lungs with TB.

**Supplementary Figure 6.** Expression profile of GBP family genes in macrophages during Mtb H37Rv infection.

**Supplementary Figure 7.** Expression of GBP1, HIF1A and inflammasome markers in GBP1 or HIF1A KD Hu-MΦ.

**Supplementary Figure 8.** Regulation of Autophagy by HIF1A and GBP1 in hu-MΦ.

**Supplementary Figure 9.** Unprocessed original images of Western blots.

## 6. Author Contributions

S.S. conceived the concept, obtained funding and supervised the studies and revised the manuscript. R.K., A.K., and G.B. designed and performed the experiments and analyzed data. S.G., S.H., and S.P. performed the microarray studies and data analysis. All authors meet the requirements for authorship in this article. All authors have read, approved and agreed for publication of this manuscript.

## 7. Acknowledgments

This study was funded in-part by a research grant from the NIAID-NIH to SS (#AI161822).

## 8. Ethics Statement

The animal studies were approved by the Rutgers University Institutional Animal Care and Use Committee (IACUC), which is compliant with the Animal Welfare Association (AWA) and United States Department of Agriculture (USDA) guidelines.

## 10. Data Availability Statement

The microarray data from rabbit and human lungs with tuberculosis were submitted to Gene Expression Omnibus (GEO) website. The accession numbers are: GSE33094; GSE39219 (rabbit lung TB) and GSE20050 (human lung TB). Other data from this study are available from the corresponding author (S.S) upon proper request.

## 13. LEGEND FOR SUPPLEMENTARY MATERIAL

Supplementary Figure 1. **Proliferation of Mtb H37Rv, HN878 and CDC1551 in rBMDM, THP-1 and hu-MΦ and differential regulation of canonical immunological pathways in rabbit lungs infected with Mtb HN878 and CDC1551.** The number of bacterial colony forming units (CFU) were measured in MtbHN878, CDC1551 or H37Rv infected rBMDM (**A**), THP-1 (**B**), and hu-MΦ (**C**) *P<0.05; **P<0.01. n=3-4 wells per group per time point and repeated twice. Statistical analyses were performed using students t test. *P<0.05; ** P< 0.01. (**D**). Heat map of canonical immunological pathways differentially regulated in the lungs of rabbits at 4 weeks after infection with HN878 or CDC1551, compared to uninfected controls. Values plotted and shown in the scale bar are z-scores. Red color indicates upregulation, and green color indicates downregulation of specific pathways based on z-score significance. Experiments were performed with n=3-4 rabbit samples in each group.

Supplementary Figure 2. **Expression of genes involved in immune pathways in the lungs of patients with TB.** Heat map of inflammatory cytokines and chemokines genes (**A**), inflammasome activation network genes (**B**), and HIF1 pathway genes (**C**) were analyzed in the lungs of TB patients with necrotic granulomas with Mtb (Hu-NG-AFB^+^) or fibrotic nodules without Mtb (Hu-FN-AFB^-^), compared to control lungs. Values plotted and shown in the scale bar are z-scores. Red color indicates upregulation, and green color indicates downregulation of specific pathways based on z-score significance. Experiments were performed with n=3-4 samples in each group.

Supplementary Figure 3. **Expressions of genes in the canonical inflammatory cytokine/chemokine signaling pathway in Mtb-infected rabbit lungs.** (**A**). Expression pattern and interaction map of inflammatory cytokine/chemokine signaling pathway genes in rabbit lungs at 4 weeks post-infection with HN878. (**B**). Heat map showing the expression pattern of inflammatory cytokine/chemokine signaling pathway genes in rabbit lungs 4 weeks post-infection with HN878 (4w-HN) and CDC1551 (4w-CDC). The red color indicates upregulation, and the green color indicates downregulation of specific genes in the pathway. Data are representatives of experiment performed on 3-4 rabbits in each group.

Supplementary Figure 4. **Expression profile of NLRP3 inflammasome activation pathway genes in macrophages during Mtb H37Rv infection.** qPCR was used to measure the transcript levels of *HIF1A, NLRP3*, *ASC*, *IL1B*, *TNFA* and *IL6* in rBMDMs (**A**), THP-1 (**B**), and hu-MΦ (**C**) infected with H37Rv for 24 or 48 hours. Fold changes in expression levels of indicated genes in Mtb-infected samples relative to those in uninfected (UI) samples were calculated. The level of *ACTNB* expression was used to normalize the expression level of test genes. Data are representative of three independent experiments performed with n=3-5 samples per group. Statistical analyses were performed using Student’s *t* -test. *P<0.05; ** P < 0.01; ***P<0.005.

Supplementary Figure 5. **Expression profile of IFN signaling pathway genes in rabbit lungs with TB.** (**A**). Pathway map showing the expression pattern and interaction of member genes in the IFN signaling pathway in HN878-infected rabbit lungs. (**B**) Heat map showing the expression pattern of inflammatory cytokine/chemokine signaling pathway genes in rabbit lungs 4 weeks after HN878 (4w-HN) and CDC1551 (4w-CDC) infection. The red color indicates upregulation, the green color indicates downregulation, and the blue color indicates no significant expression of genes. Data are representatives of experiment performed on 3-4 rabbits in each group.

Supplementary Figure 6. **Expression profile of GBP family genes in macrophages during Mtb H37Rv infection.** (A-C) Transcript levels of *GBP1*, *GBP2*, *GBP3*, *GBP4* and *GBP5* in rBMDM (**A**), THP-1 (**B**), and hu-MΦ (**C**) infected with MtbH37Rv for 24 or 48 hours measured by qPCR. Fold changes of gene expression levels in Mtb-infected samples relative to uninfected samples were calculated. The level of *ACTNB* expression was used to normalize the expression level of test genes. Experiments were repeated thrice with n=3-4 samples and statistical analyses were performed using the student *t*-test. *P<0.05; ** P < 0.01; ***P<0.005; ****P<0.001.

Supplementary Figure 7. **Expression of GBP1, HIF1A and inflammasome markers in GBP1 or HIF1A KD cells.** The THP-1 macrophages were treated with siRNAs against *HIF1A* (top panels) or *GBP1* (bottom panels) or scrambled siRNA (Cont) and the mRNA level of *GBP1*, *HIF1A*, *NLRP3*, *ASC* and *IL1B* was determined by qRT-PCR. Expression level of each gene in the control cells is set to 1 and the expression level in siRNAs transfected and Mtb-infected samples is expressed as relative fold change. The expression level of *ACTNB* in each sample was used to normalize the expression of test genes in respective samples. Experiments were repeated thrice with n=3-4 samples and statistical analyses were performed using the student *t*-test. *P<0.05; ** P < 0.01; ***P<0.005; ****P<0.001.

Supplementary Figure 8. **Regulation of Autophagy by HIF1A and GBP1 in macrophages.** THP-1 macrophages were treated with specific siRNAs against *GBP1* or *HIF1A* or with scrambled siRNA (Control), infected with CDC1551 or HN878 or remain uninfected, then stained with an antibody against the autophagic marker LC3 (red color). DAPI was used to stain nuclei (blue color). Fluorescence-based immunocytochemistry was performed on the macrophages to visualize specific stains. The scale bar is 50 microns and applies to all panels. The images are representative of experiments performed thrice with two technical replicates.

Supplementary Figure 9. **Unprocessed original images of Western blots.** Raw, unprocessed original western blot images of various target proteins at low and high exposures. The portion of the images used as a composite in Figure 4 is marked with red boxes in this image. The images are representative of experiments performed thrice with two technical replicates.

