## Supplementary figures and images for "Inflammasome activation differences underpin different *Mycobacterium tuberculosis* infection outcomes"

### Supplemental Files_Combined

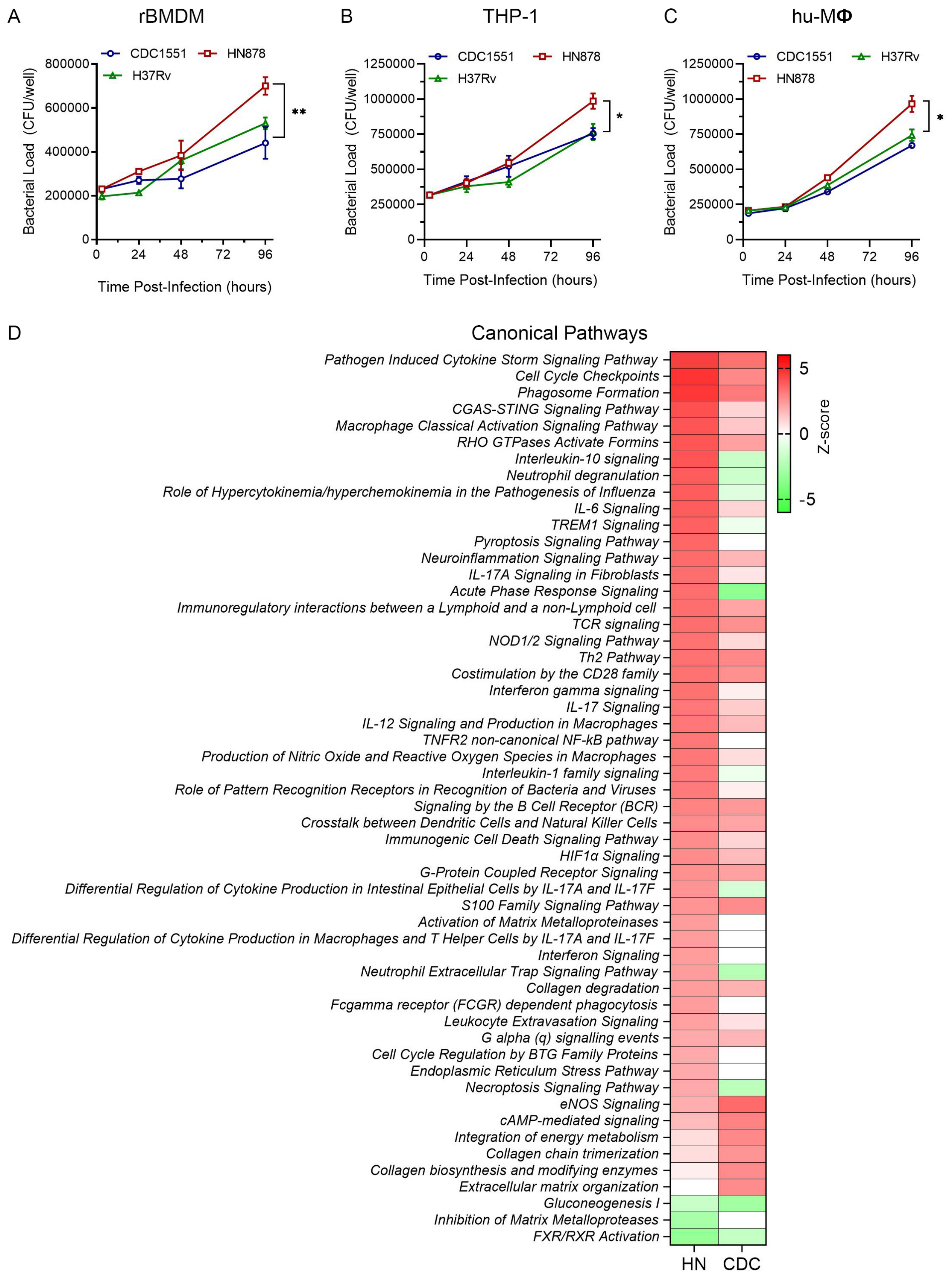

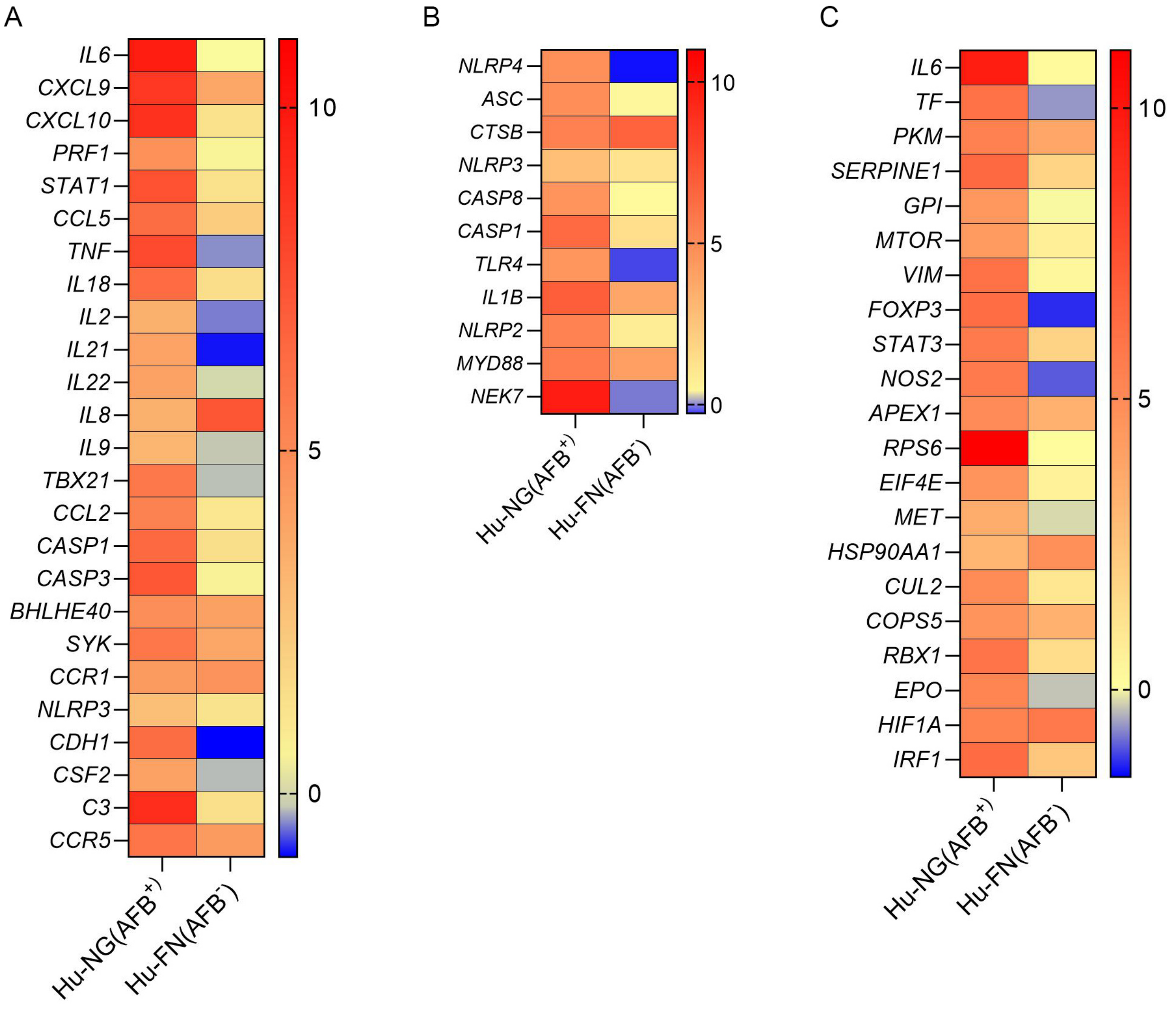

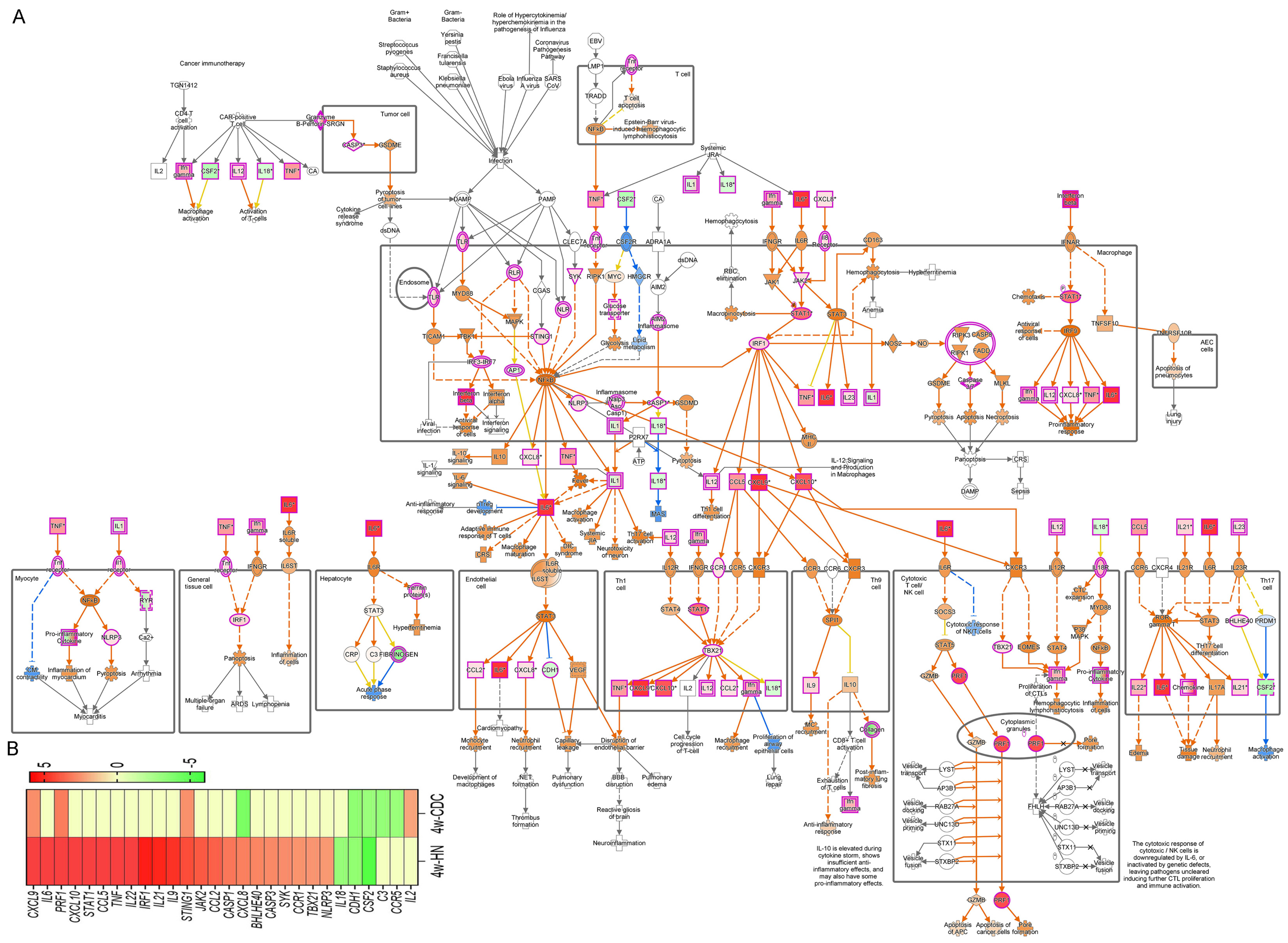

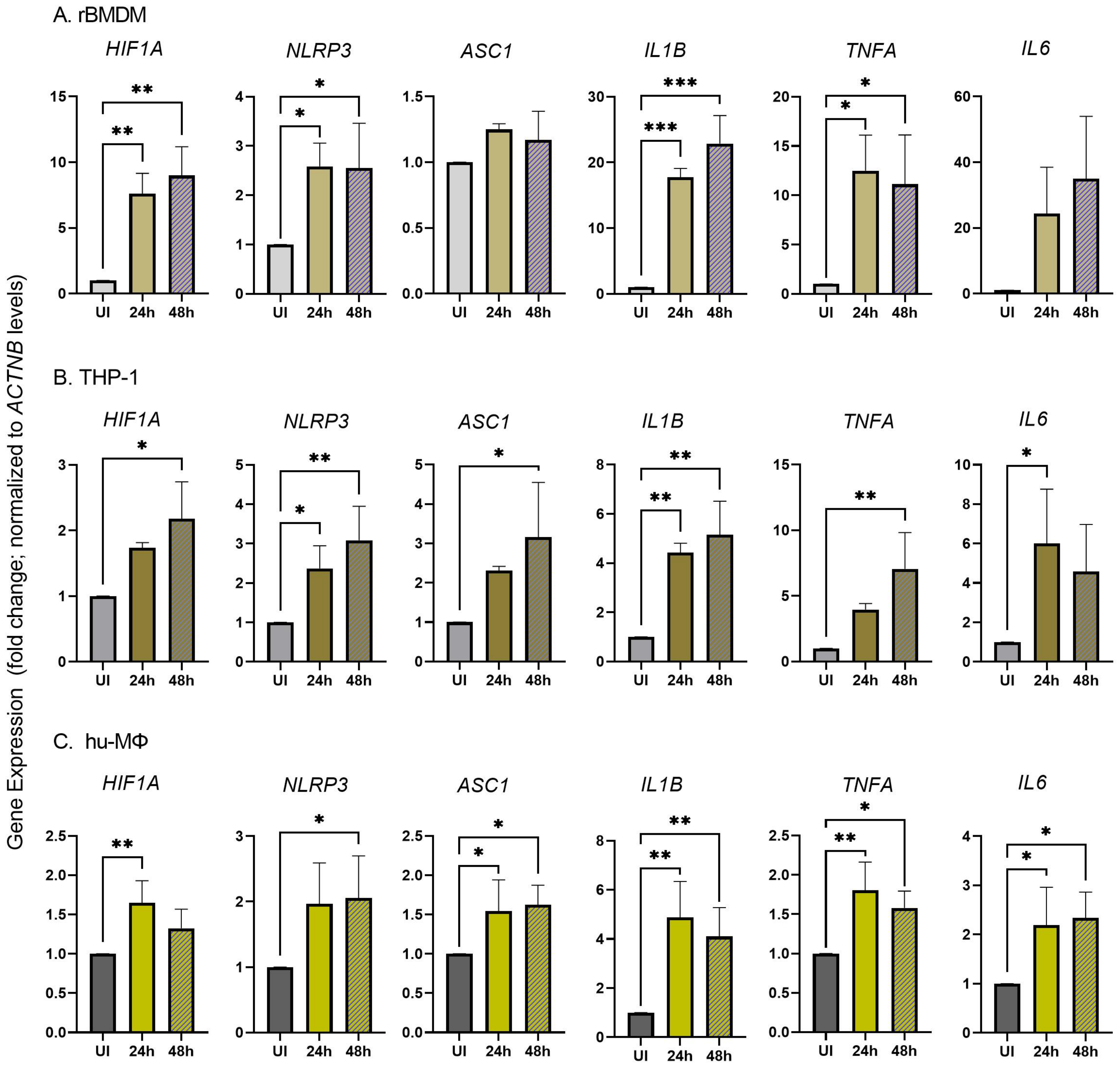

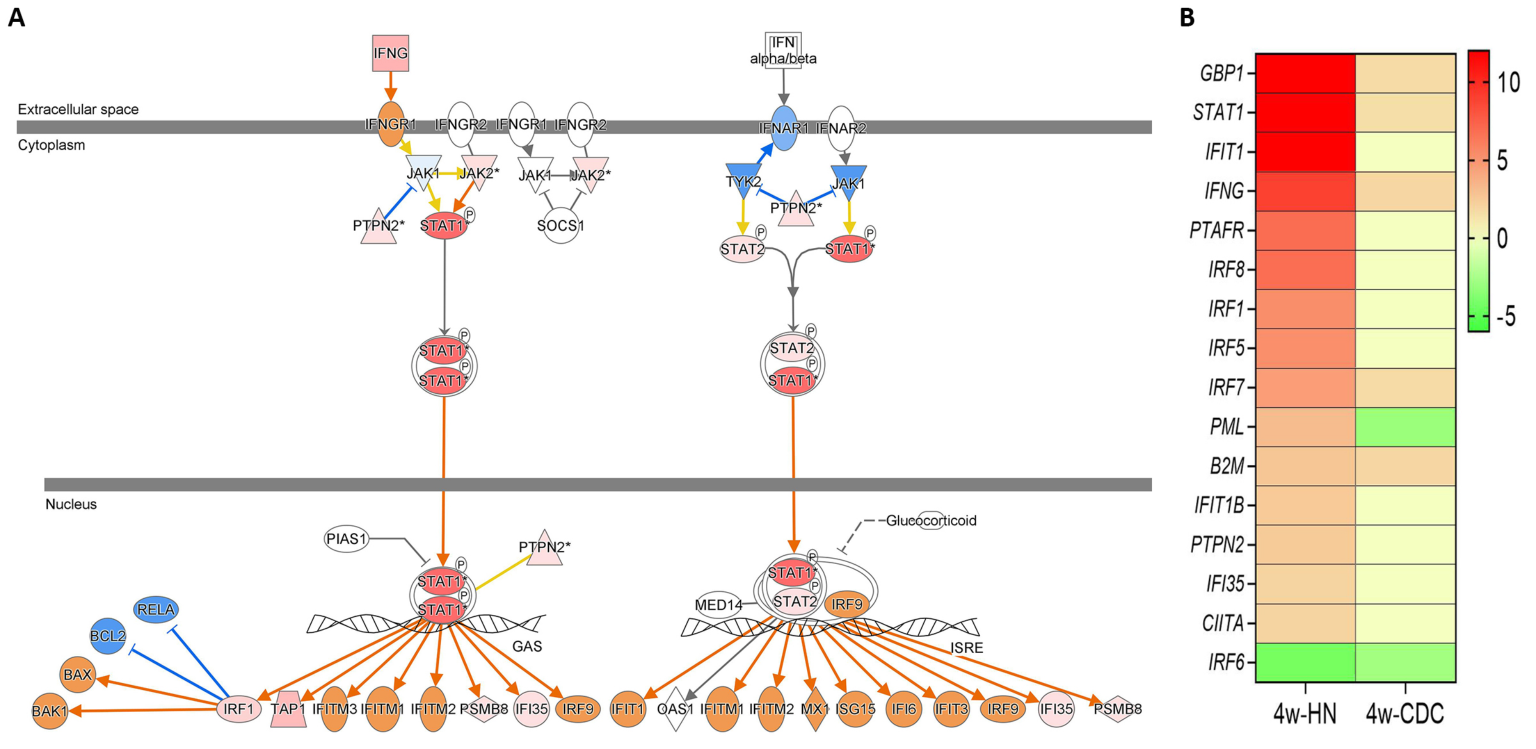

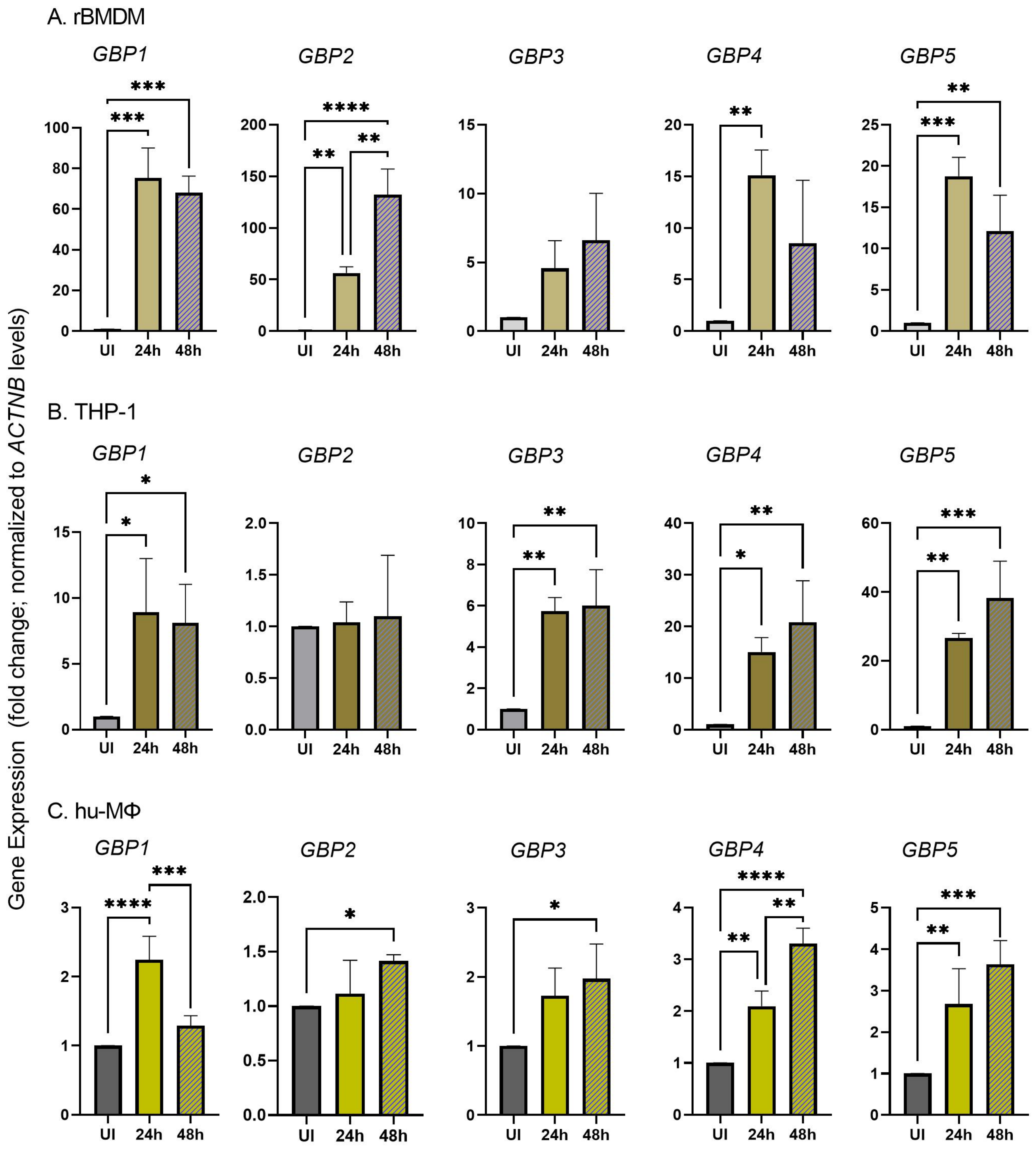

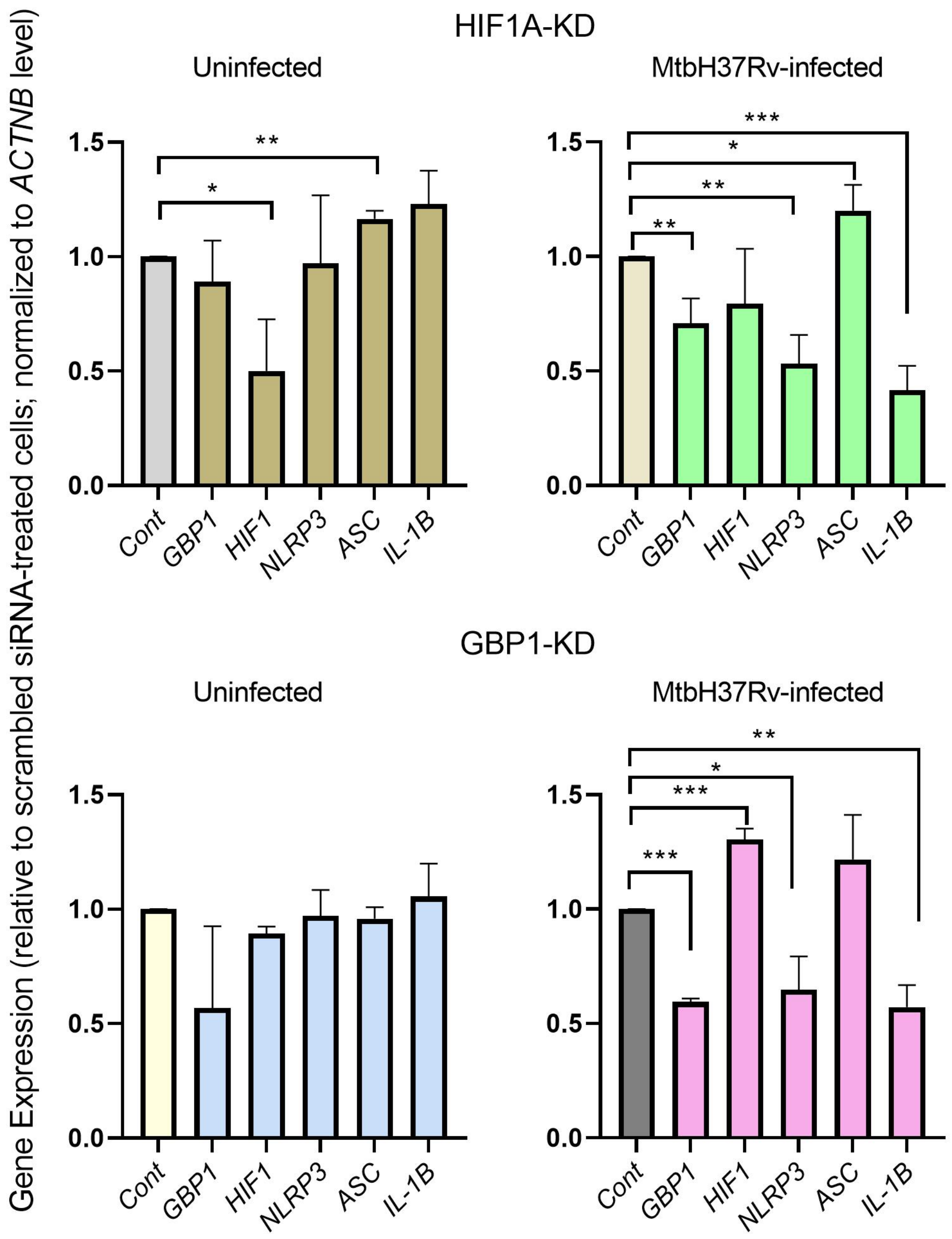

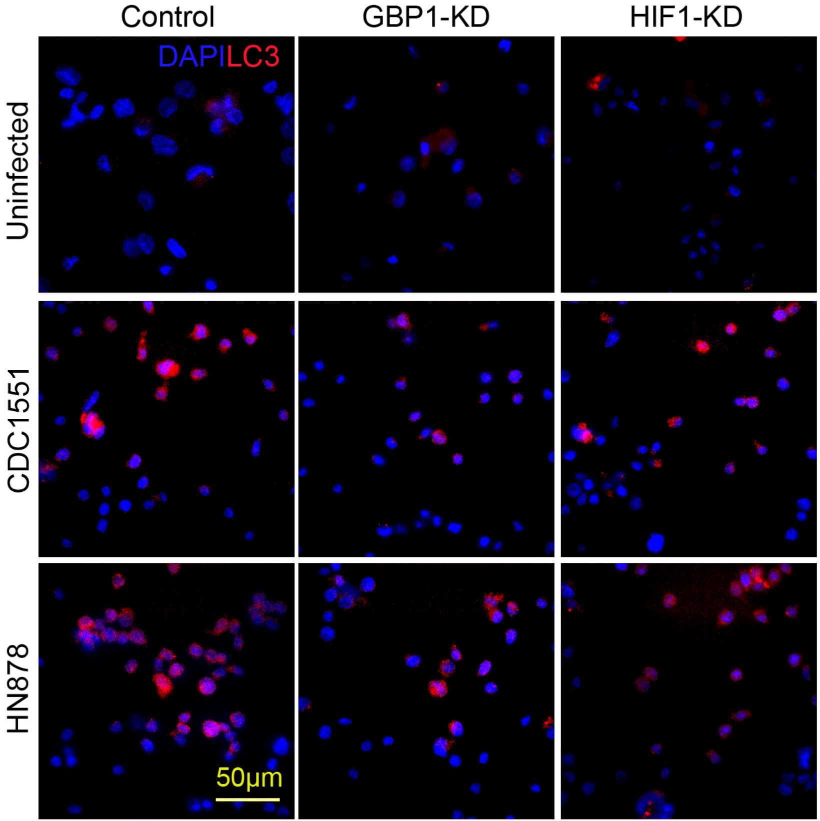

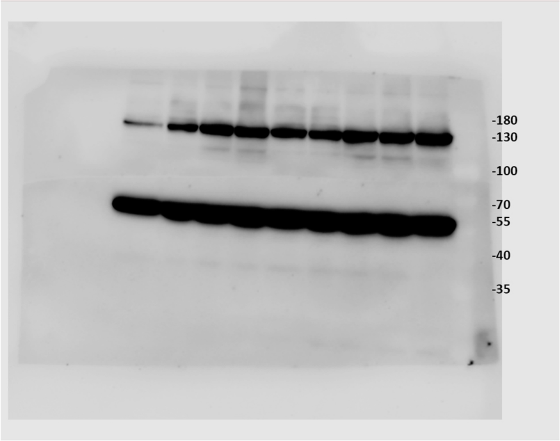


High Exposure

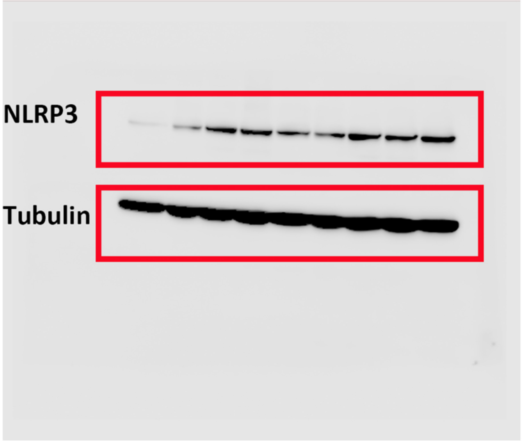


Low Exposure

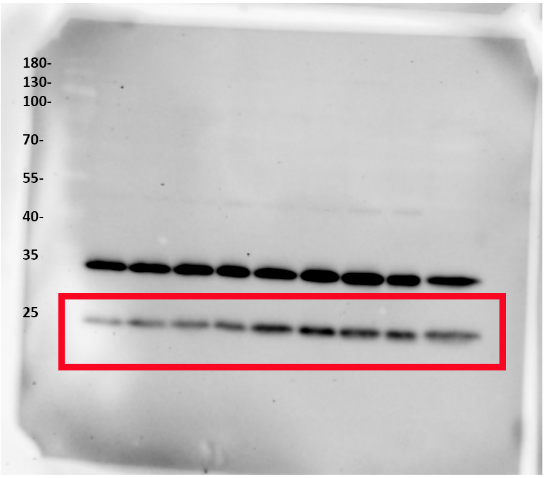


ASC

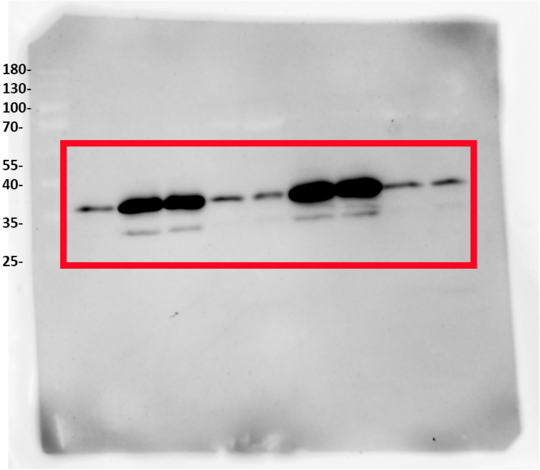


IL1B

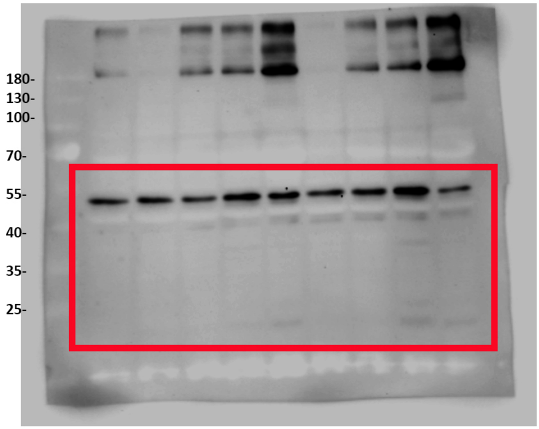


Caspase

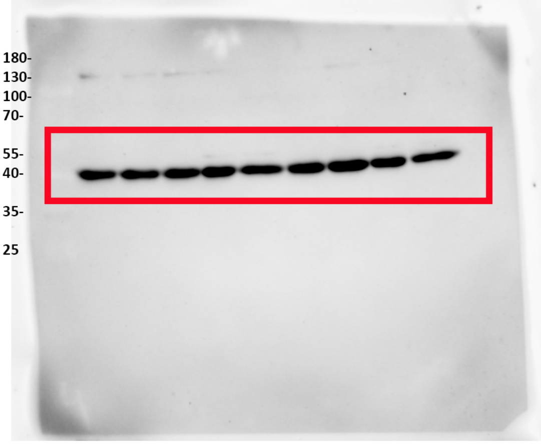


ACTIN
